# The gut microbiota directs vitamin A flux to regulate intestinal T cell development

**DOI:** 10.1101/2025.09.08.674524

**Authors:** Tarun Srinivasan, Chaitanya Dende, Kelly A. Ruhn, Stefanie L. Moye, Ann Johnson, Cassie L. Behrendt, Brian Hassell, Gonçalo Vale, Jiwoong Kim, Jake N. Lichterman, Chaoying Liang, Carlos Arana, Prithvi Raj, Xiaowei Zhan, Jeffrey G. McDonald, Andrew Y. Koh, Lora V. Hooper

**Affiliations:** Department of Immunology, University of Texas Southwestern Medical Center, Dallas, TX, USA; Department of Surgery, University of Texas Southwestern Medical Center, Dallas, TX, USA; Center for Human Nutrition, University of Texas Southwestern Medical Center, Dallas, TX, USA; Department of Internal Medicine, University of Texas Southwestern Medical Center, Dallas, TX, USA; Department of Public Health, O’Donnell School of Public Health, University of Texas Southwestern Medical Center, Dallas, TX, USA; Department of Molecular Genetics, University of Texas Southwestern Medical Center, Dallas, TX, USA; Department of Pediatrics, University of Texas Southwestern Medical Center, Dallas, TX, USA; Harold C. Simmons Comprehensive Cancer Center, University of Texas Southwestern Medical Center, Dallas, TX, USA; Howard Hughes Medical Institute, University of Texas Southwestern Medical Center, Dallas, TX, USA

**Keywords:** microbiota, vitamin A, retinoids, serum amyloid A, CD4⁺ T cells, mesenteric lymph nodes, myeloid cells, retinoic acid receptor (RAR), intestinal homing, postnatal development

## Abstract

The intestinal microbiota shapes adaptive immunity, but the mechanisms remain incompletely defined. Here, we show that the microbiota initiates the movement of retinoids—dietary vitamin A derivatives including retinol and retinoic acid—through a sequential pathway from epithelial cells to myeloid cells and ultimately to T cells in the mesenteric lymph nodes (mLNs). This cellular axis is traversed over three days. Microbe-associated molecular patterns (MAMPs) initiate retinoid flux by inducing expression of serum amyloid A (SAA) proteins. These epithelial retinol-binding proteins are necessary and sufficient for epithelial-to-myeloid cell retinoid transfer and for myeloid cell migration to the mLNs. In the mLNs, microbial antigen drives retinoid transfer from myeloid cells to developing T cells, culminating in T cell retinoid uptake and transcriptional programming. This pathway is activated during postnatal development, when gut adaptive immunity is first established. These findings reveal that the microbiota programs intestinal adaptive immunity by regulating immune cell access to a nutrient-derived developmental signal.

**Highlights:**

- The gut microbiota enables vitamin A flux to developing intestinal CD4⁺ T cells.
- Microbiota-induced SAA initiates vitamin A flux along a gut myeloid–T cell axis.
- Microbial molecular patterns and antigen drive distinct steps of vitamin A flux.
- Microbiota-driven vitamin A flux programs intestinal T cell homing and maturation.

**Graphical Abstract:** 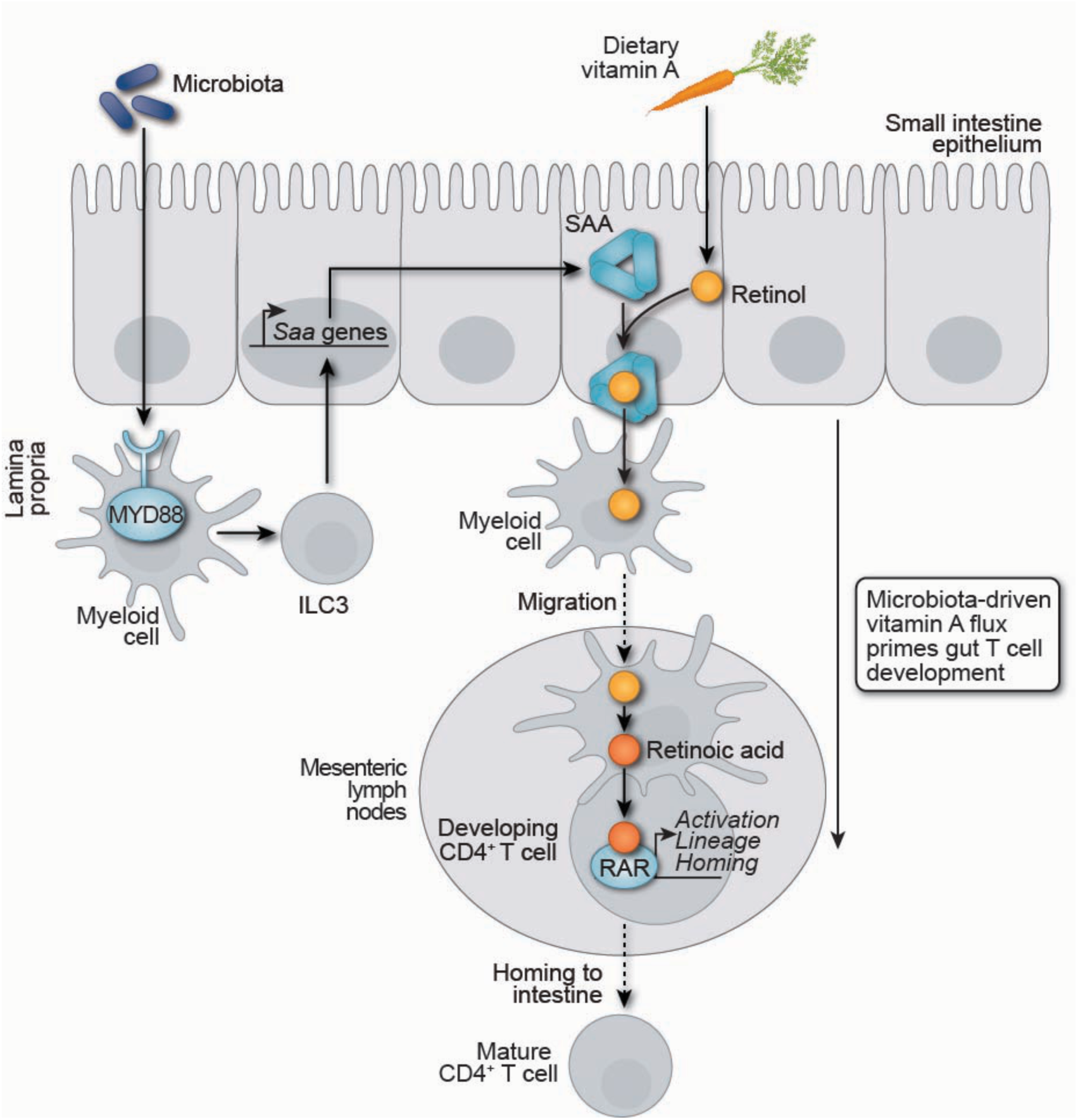

## Introduction

Beginning at birth, the mammalian intestine is colonized by a dense and diverse community of microorganisms.^1,2^ These organisms, primarily bacteria, form a symbiotic relationship with their host, enhancing digestion, promoting nutrient absorption, and providing resistance to pathogen colonization.^3–6^ Importantly, resident gut microbes are also required for normal development of the intestinal adaptive immune system.^7^ Despite this established role, the molecular mechanisms by which the microbiota instructs adaptive immune cell development remain incompletely understood.

The intestinal adaptive immune system is comprised of T and B lymphocytes that restrict microbial entry into tissues and contain organisms that breach the epithelial barrier. Studies in germ-free mice demonstrate that, without the microbiota, the intestine is sparsely populated with lymphocytes and exhibits impaired immune maturation.^7–9^ Microbial colonization restores intestinal lymphocyte accumulation, highlighting the essential role of the microbiota in shaping intestinal immunity.^10–12^

CD4^+^ T cells illustrate the dependence of gut adaptive immune cell development on microbial cues. These cells are first primed in the mesenteric lymph nodes (mLNs), secondary lymph nodes located outside the intestinal wall.^13,14^ Central T cell development in the thymus establishes a naïve repertoire, whereas peripheral development in the mLNs and intestine reflects programming and maturation in response to microbial and tissue-derived signals.^15^ Thus, signals originating at the intestinal surface or in the lamina propria must traverse anatomically distinct tissue compartments to reach the mLNs and promote T cell maturation and intestinal homing.

Microbial antigen is one such signal that moves from the lumen, through the lamina propria, and to the mesenteric lymph nodes to prime adaptive immunity. Antigen-presenting myeloid cells in the lamina propria capture microbial antigens, migrate to the mLNs, and prime naïve CD4^+^ T cells.^13,14,16–18^ However, antigen alone is insufficient to program intestinal T cell maturation and homing. In vitamin A-deficient mice, T cell migration to the intestine is impaired despite the presence of a microbiota, indicating that, in addition to microbial antigen, a vitamin A-derived signal is needed.^19,20^

Vitamin A enters the diet as β-carotene and retinyl esters, which are absorbed by the intestinal epithelium and metabolized to retinol.^21,22^ Retinol belongs to the broader class of compounds known as retinoids, which are vitamin A-derived molecules with diverse metabolic and signaling functions.^23,24^ A key retinoid, retinoic acid (RA), is generated from retinol by intestinal myeloid cells and transferred into developing mLN T cells, directing them to the intestine.^25–30^ RA does this by activating T cell retinoic acid receptors (RARs), thereby inducing expression of gut-homing receptors which guide migration to the lamina propria.^19^ Together with data from germ-free mice, these findings show that both vitamin A and the microbiota are essential drivers of T cell development. Yet how RA and microbiota-derived cues integrate to regulate T cell maturation and intestinal homing remains unclear.

Here, we show that the microbiota directs vitamin A-derived retinoids to developing mLN T cells to enable their maturation and homing via activation of RARs. Microbial colonization initiates a stepwise transfer of retinoids from intestinal epithelial cells to myeloid cells and ultimately to developing T cells in the mLNs. This epithelial– myeloid–T cell axis is traversed over three days and activates a transcriptional program that enables T cell maturation and migration to the small intestine. The process begins when microbial colonization, acting through innate immune sensing pathways, induces expression of serum amyloid A (SAA) proteins—retinol-binding proteins that deliver retinol from the epithelium to intestinal myeloid cells.^31^ SAA is sufficient, even in the absence of the microbiota, to drive retinoid loading of lamina propria myeloid cells and their migration to the mLNs. Antigen, however, is required for the transfer of retinoids from mLN myeloid cells to T cells. Notably, microbiota-dependent retinoid flux occurs during early postnatal development, a critical window for adaptive immune development that coincides with maturation of the gut microbiota. Together, these findings define a mechanism by which the microbiota communicates a developmental signal across tissue compartments to mLN T cells and reveal how microbial and dietary cues converge to regulate intestinal adaptive immunity.

## Results

### The microbiota directs vitamin A flux along an epithelial–myeloid–T cell axis

RA is a key developmental signal for intestinal T cells that undergo maturation in the mLNs, which lie outside the gut wall.^19,20^ Because RA is generated from dietary vitamin A, it must traverse a cellular axis to travel from the gut lumen to the mLNs. Vitamin A is first absorbed by intestinal epithelial cells (IECs), where it is converted to retinol.^22^ Specialized CD11c⁺ myeloid cells in the lamina propria (LP), the immune-rich connective tissue beneath the intestinal epithelium, acquire retinol from IECs.^31^ These cells convert retinol to RA through sequential oxidation by retinol dehydrogenase and retinaldehyde dehydrogenase, then migrate to the mLNs to deliver RA to naïve CD4⁺ T cells.^25–29^ This RA induces the expression of homing receptors such as CCR9, enabling mature T cells to exit the mLNs and populate the intestinal LP as functional T cells (Fig. 1A).^19^

**Figure 1:**
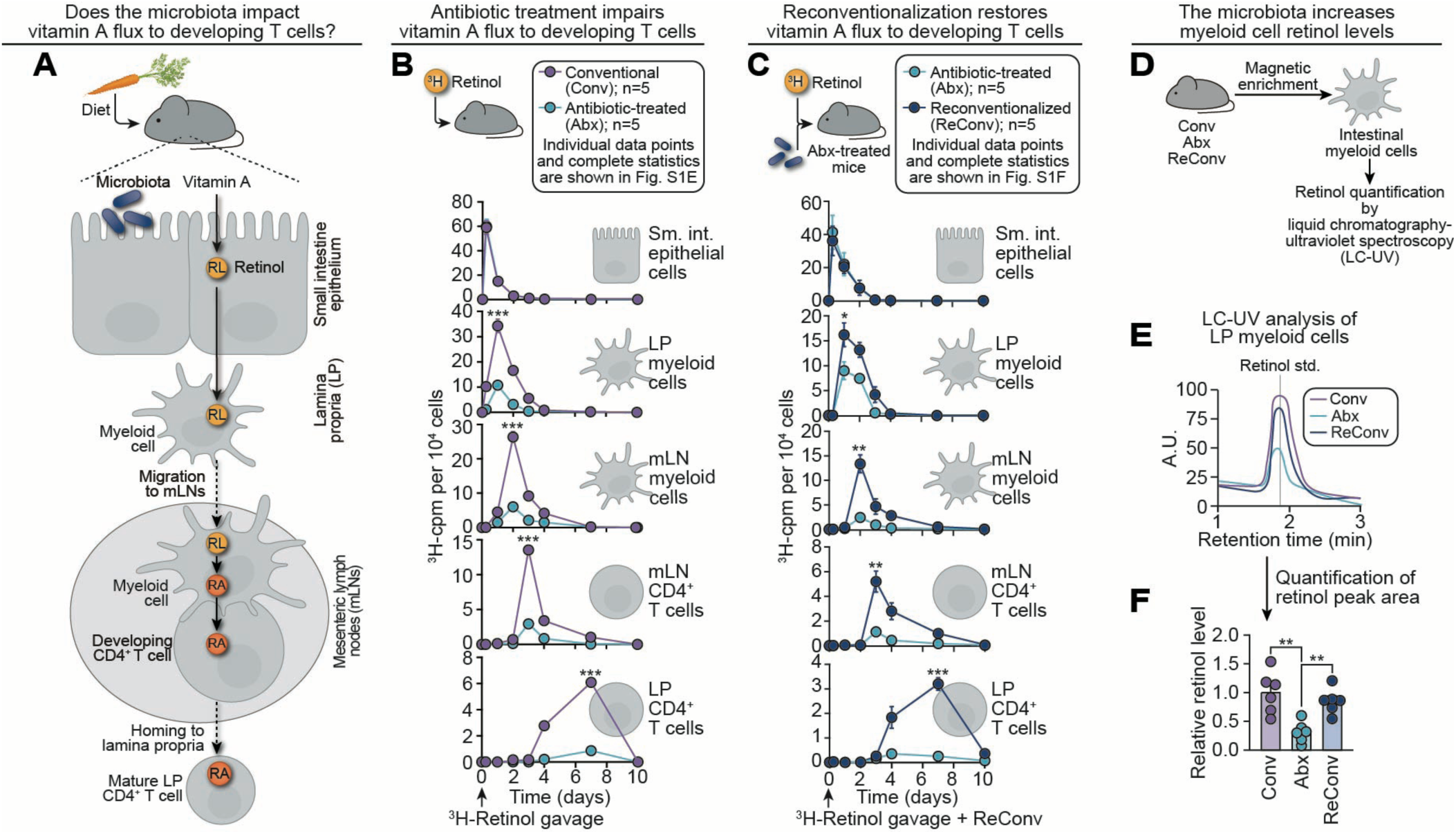
The microbiota directs vitamin A flux along an epithelial–myeloid–T cell axis. **(A)** Dietary vitamin A is absorbed by intestinal epithelial cells (IECs) and converted to retinol (RL).^22^ Retinol is transferred to lamina propria (LP) myeloid cells, which migrate to the mesenteric lymph nodes (mLNs).^16,31^ The myeloid cells convert retinol to retinoic acid (RA) and deliver it to naïve CD4⁺ T cells.^25–29^ RA is a critical developmental signal that promotes T cell maturation and homing to the lamina propria of the intestine.^19^ How the microbiota impacts the delivery of this diet-derived developmental signal to developing T cells is unknown. **(B)** Movement of retinoids (vitamin A-derived compounds) through the IEC–myeloid cell–T cell axis over time (flux) in conventional and antibiotic-treated mice. ^3^H-retinol (1 μCi) was administered to conventional (Conv) and antibiotic-treated (Abx) mice by gavage. IECs were isolated using EDTA; LP CD11c⁺ myeloid cells, mLN CD11c⁺ myeloid cells, mLN CD4⁺ T cells, and LP CD4⁺ T cells were isolated by magnetic enrichment. Radioactivity (cpm) was measured at 0 and 6 hours, and 1, 2, 3, 4, 7, and 10 days post-gavage and normalized to cell number. n=5 mice per group per time point. Individual data points and complete statistics are in Figure S1E. **(C)** Movement of retinoids through the IEC–myeloid cell–T cell axis in antibiotic-treated and reconventionalized (ReConv) mice after antibiotic treatment. ³H-retinol was administered to mice alongside reintroduction of a conventional microbiota after antibiotic treatment. ^3^H-cpm were measured in isolated cells as in (B). n=5 mice per group per time point. Individual data points and complete statistics are in Figure S1F. **(D)** Retinol was measured in isolated LP CD11c⁺ myeloid cells by liquid chromatography ultraviolet spectroscopy (LC-UV). **(E)** Representative LC-UV chromatograms of retinol detected in LP CD11c⁺ cells isolated from the small intestines of conventional, antibiotic-treated, and reconventionalized mice. Samples were normalized for protein prior to loading. Retinol was identified by comparison to a purified standard, which eluted at ∼1.9 minutes, and quantified by UV absorbance at its absorbance maximum (λₘₐₓ = 325 nm). **(F)** Quantification of retinol levels by LC-UV, normalized to protein. The average of the values in conventional mice was set to 1; antibiotic-treated and reconventionalized values were expressed relative to this average. Each point represents one mouse. RL, retinol; RA, retinoic acid; LP, lamina propria, mLN, mesenteric lymph node; Conv, conventional, Abx, antibiotic-treated; ReConv, reconventionalized; cpm, counts per minute; LC-UV, liquid chromatography–ultraviolet spectroscopy; A.U., absorbance units; min, minutes. Data are representative of at least two independent experiments. Means ± SEM are plotted. \**p* < 0.05; \*\**p* < 0.01; \*\*\**p* < 0.001 by two-tailed Student’s *t*-test. See also Figures S1 and S2.

Because the microbiota promotes intestinal T cell development,^5,32–34^ we hypothesized that it might do so in part by regulating the mobilization of dietary retinoids along this cellular axis. To test this, we gavaged conventional and antibiotic-treated mice with radiolabeled retinol (³H-retinol) and tracked radioactivity in IECs, CD11c⁺ myeloid cells, and CD4⁺ T cells over 10 days. These cell populations were purified by EDTA treatment (IECs) or magnetic isolation (myeloid and T cells), with enrichment validated by flow cytometry (Fig. S1A–D). Retinoid counts were normalized to cell number to control for antibiotic-induced shifts in immune cell frequency (Fig. S2A–G).

In conventional mice, ³H-retinoids moved sequentially from IECs to LP CD11c⁺ myeloid cells, then to mLN CD11c⁺ myeloid cells, mLN CD4⁺ T cells, and finally LP CD4⁺ T cells (Fig. 1B; Fig. S1E). Although retinoid absorption by IECs was similar in conventional and antibiotic-treated mice, downstream flux through LP myeloid cells and subsequently to mLN myeloid cells and CD4⁺ T cells was markedly impaired in antibiotic-treated mice. Notably, systemic retinoid levels in serum and liver remained unchanged (Fig. S1G), indicating that the microbiota selectively influences retinoid flux through intestinal cellular pathways without altering whole-body retinoid trafficking and distribution. Reconventionalization–the reintroduction of the intestinal microbiota after antibiotic treatment–rescued ³H-retinoid movement through the IEC–myeloid–T cell axis, supporting microbiota-dependent regulation of retinoid flux (Fig. 1C; Fig. S1F).

We next quantified the proportion of absorbed retinoids allocated to the myeloid–T cell axis. ³H-retinoid distribution was measured in IECs, LP CD11c⁺ cells, serum, and liver at 6, 12, and 24 hours after gavage. At 24 hours, ∼57% of ³H-retinoids had localized to the liver, ∼3% to serum, ∼22% remained in IECs, and <2% was detectable in LP myeloid cells (Fig. S1H). These data show that, while most dietary retinoids traffic to the liver, a small but significant fraction is routed to the intestinal myeloid–T cell axis.

To assess the chemical identity of retinoids within LP CD11c^+^ myeloid cells, we quantified retinol levels in these cells using liquid chromatography–ultraviolet (LC-UV) spectroscopy. Consistent with the radioactive tracer data, antibiotic-treated mice had reduced retinol levels in LP myeloid cells compared to both conventional and reconventionalized mice (Fig. 1D–F). As expected, we did not detect retinol in CD4⁺ T cells, since T cells acquire RA from myeloid cells rather than generating it from retinol.^19,25^ We were unable to measure RA due to its lability and rapid turnover, which limit cellular detection by LC-UV. Taken together, these data indicate that the microbiota initiates retinoid flux through an IEC–myeloid–T cell axis in the small intestine.

### The microbiota directs RAR activation along an intestinal myeloid–T cell axis

RA, the bioactive metabolite of vitamin A, binds RARs to drive gene transcription.^35,36^ Because the microbiota regulates retinoid movement through the intestinal immune system, we hypothesized that it also directs the sequential activation of RAR signaling through the myeloid–T cell axis. To test this, we employed RARE*-lacZ* reporter mice, in which β-galactosidase (LacZ) is expressed under the control of three tandem RAR response elements in all cells.^37^ Following antibiotic treatment and microbial reconstitution, we quantified RAR activity in LP and mLN CD11c⁺ myeloid cells and CD4⁺ T cells over 10 days. RAR activity was measured by flow cytometry using fluorescein di-V-galactoside (FDG), a β-galactosidase substrate metabolized by LacZ-expressing cells to yield fluorescein, which is detectable by flow cytometry (Fig. 2A). We observed a stepwise induction of RAR activity: first in LP CD11c⁺ myeloid cells, then in mLN CD11c⁺ myeloid cells, followed by mLN CD4⁺ T cells, and finally LP CD4⁺ T cells (Fig. 2B,C; Fig. S3A). This sequence mirrors the pattern of retinoid transfer, indicating that transit of retinoids across this axis coincides with the activation of RAR signaling.

**Figure 2:**
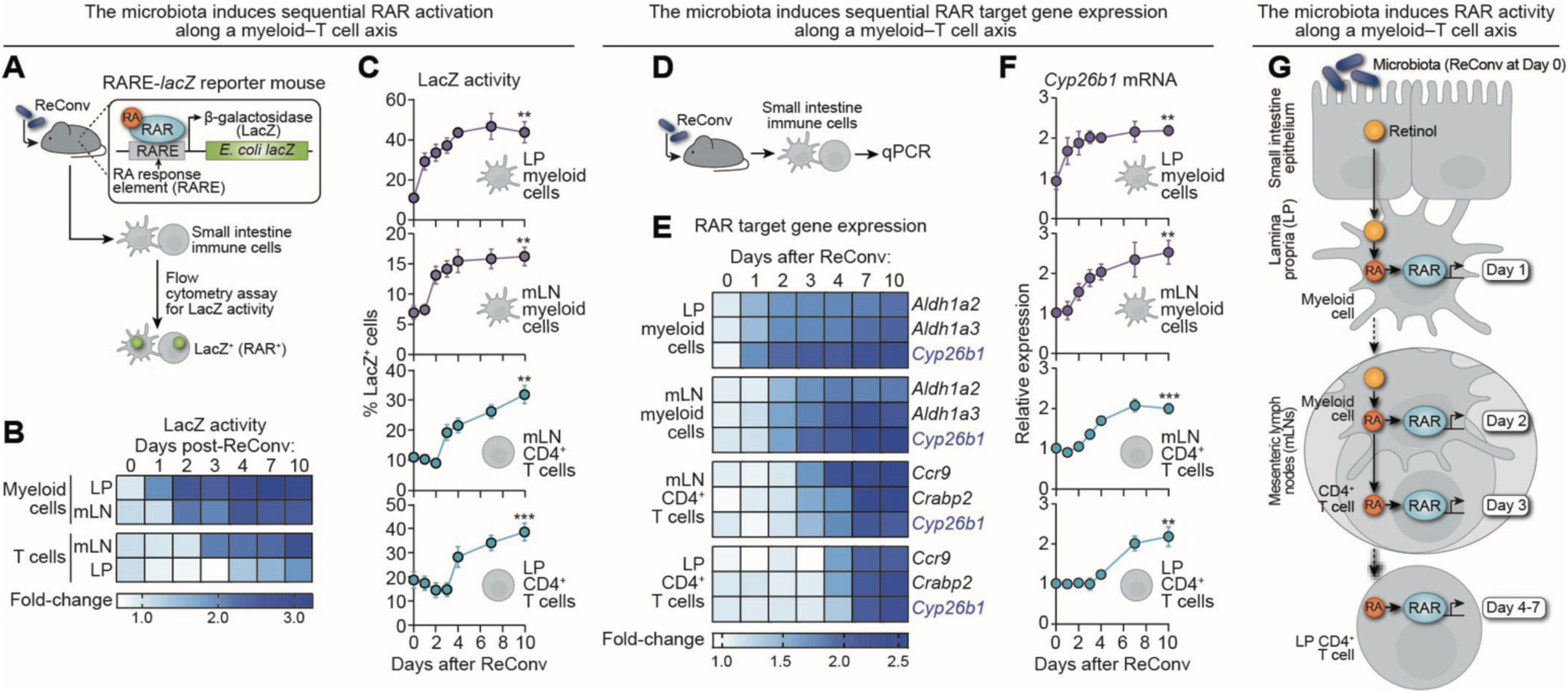
The microbiota directs RAR activation along an intestinal myeloid–T cell axis. **(A)** Strategy for assessing RAR activity in response to microbial colonization. RARE*-lacZ* reporter mice harbor a transgene that drives β-galactosidase (LacZ) expression under the control of three tandem retinoic acid response elements (RAREs).^37^ RARE-*lacZ* mice were treated with antibiotics for 10 days and then reconventionalized. LacZ activity was quantified by flow cytometry using fluorescein di-V-galactoside (FDG), which fluoresces upon LacZ-mediated cleavage. RAR^+^ cells were defined by high LacZ activity. **(B)** Heat map depicting fold change in the frequency of RAR⁺ (LacZ⁺) cells over time, relative to day 0, across the myeloid–T cell axis following reconventionalization. **(C)** Frequency of RAR⁺ cells across the myeloid–T cell axis after reconventionalization. n=5 mice per group per time point. **(D)** Strategy for assessing microbial regulation of RAR target gene expression. Wild-type mice were treated with antibiotics and reconventionalized, and qPCR for RAR target genes was performed on intestinal immune cells isolated by magnetic sorting. **(E)** Time course of RAR target gene expression in CD11c^+^ myeloid cells and CD4⁺ T cells in the LP and mLNs. **(F)** *Cyp26b1* expression kinetics across each of the four cell populations following microbial reconstitution. n=5 mice per group per time point. Additional RAR target genes are shown in Fig. S3F. **(G)** Graphical model showing microbiota-directed activation of RARs along the myeloid–T cell axis over time. ReConv, reconventionalized; RA, retinoic acid; RAR, retinoic acid receptor; RARE, retinoic acid response element; LacZ, β-galactosidase. Data are representative of at least two independent experiments. Means ± SEM are plotted. \*\**p* < 0.01; \*\*\**p* < 0.001 by two-tailed Student’s *t*-test. See also Figure S3.

Importantly, flow cytometry and immunofluorescence imaging revealed that, while antibiotic-treated RARE*-lacZ* mice had markedly reduced RAR activity in the lamina propria, RAR activation in IECs was unaffected (Fig. S3B–E). This is consistent with our finding that the microbiota does not regulate epithelial retinoid uptake but rather its mobilization to downstream immune cells.

To determine whether microbial regulation of RAR activity initiates gene expression programs along the myeloid–T cell axis, we isolated CD11c⁺ myeloid cells and CD4⁺ T cells from the lamina propria and mLNs following microbial reconstitution and measured expression of known RAR target genes (Fig. 2D). In myeloid cells, we assessed expression of *Aldh1a2* and *Aldh1a3* (RA synthesis) and *Cyp26b1* (RA degradation).^38,39^ In CD4⁺ T cells, we quantified *Ccr9* (homing receptor), *Crabp2* (RA chaperone), and *Cyp26b1* expression.^19,39,40^ As with RAR activity, target gene expression followed a stepwise induction pattern along the myeloid–T cell axis (Fig. 2E; Fig. S3F), with *Cyp26b1* expression serving as a marker of progressive RAR activation across all cell types (Fig. 2F).

Together, these findings demonstrate that the microbiota directs retinoid flux, orchestrates RAR activation, and promotes RA-dependent transcriptional programming along a myeloid–T cell axis in the small intestine (Fig. 2G).

### The microbiota initiates intestinal vitamin A flux by inducing serum amyloid A expression

We next investigated the mechanism by which the microbiota initiates dietary vitamin A flux to intestinal immune cells. We previously identified the serum amyloid A (SAA) family of retinol-binding proteins (SAA1–4).^41^ Of these, SAA1–3 are expressed in the intestine, with SAA1/2 as dominant and largely redundant paralogs.^31,42^ Produced by IECs, SAAs are microbiota-inducible and deliver retinol to intestinal myeloid cells through an endocytic cell-surface receptor.^31,41,42^ Building on these findings, we assessed the role of SAAs in facilitating microbiota-dependent intestinal retinoid flux.

Antibiotic treatment reduced intestinal *Saa1* expression, confirming previous findings (Fig. 3A),^31,41^ while expression of genes encoding the retinol-binding proteins *Rbp1* and *Rbp4* did not decrease (Fig. S4A).^43,44^ To directly assess the role of SAA in regulating intestinal retinoid flux, we used *Saa⁻^/^⁻* mice lacking all *Saa* isoforms (*Saa1–4*).^31,42^ ^3^H-retinoid tracking revealed reduced retinoid levels in lamina propria CD11c⁺ myeloid cells from *Saa⁻^/^⁻* mice one day after gavage, phenocopying microbiota-depleted controls (Fig. 3B,C). Importantly, IEC, serum, and liver ^3^H-retinoid levels were unchanged by the presence or absence of SAA, suggesting that SAA expression does not affect retinoid absorption from the diet or its systemic distribution (Fig. S4B). LC-UV analysis confirmed decreased retinol content in lamina propria CD11c^+^ myeloid cells from *Saa⁻^/^⁻* mice and antibiotic-treated controls (Fig. 3D,E).

**Figure 3:**
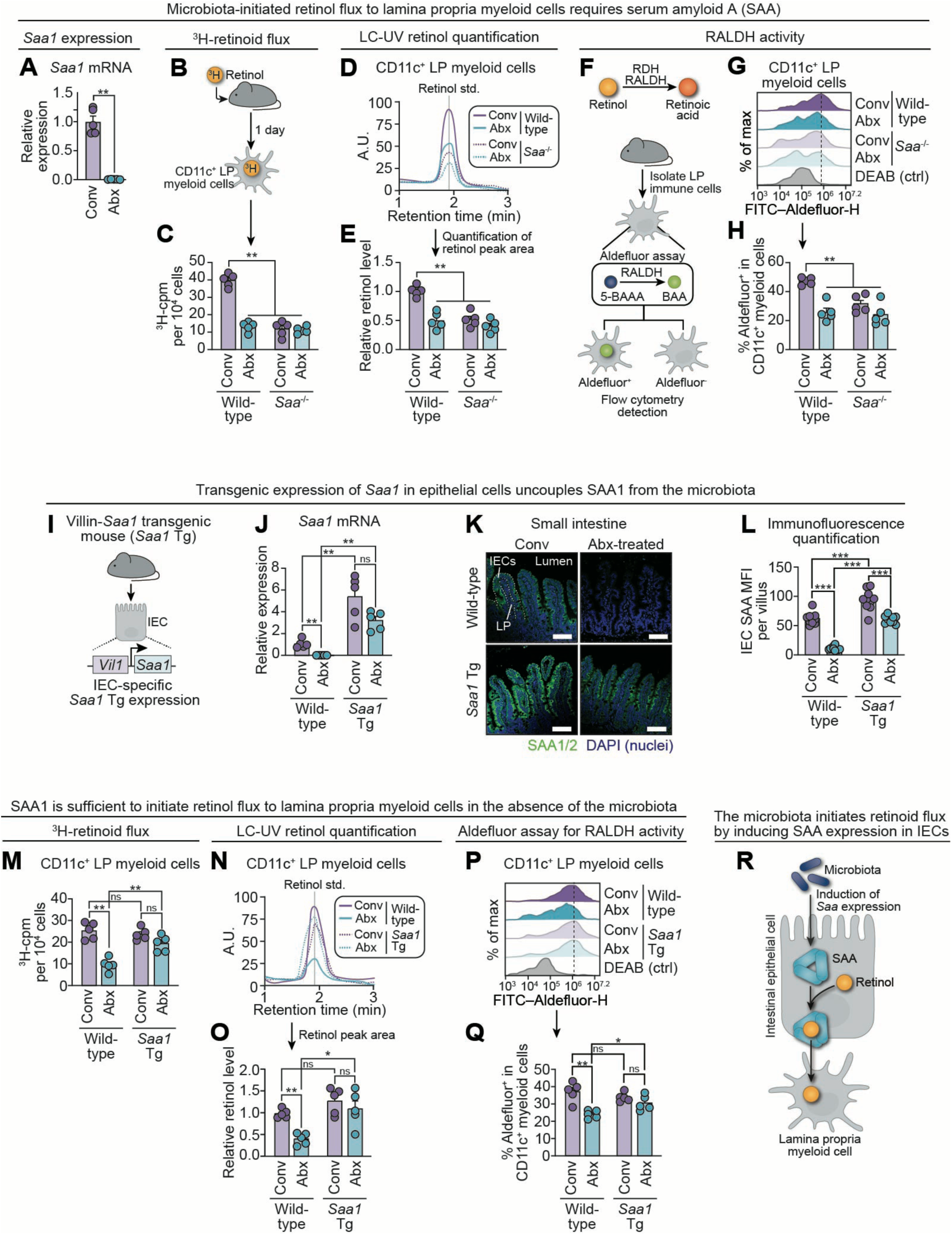
The microbiota initiates intestinal vitamin A flux by inducing serum amyloid A expression. **(A)** Small intestinal *Saa1* expression, measured by qPCR, in conventional and antibiotic-treated mice. *Saa1* encodes a microbiota-induced retinol-binding protein expressed in IECs. **(B)** ^3^H-retinol was administered to mice by gavage. LP myeloid cells were isolated by magnetic sorting 1 day post-gavage, and ^3^H-cpm were quantified by scintillation counting. **(C)** ³H-retinoids in lamina propria CD11c⁺ myeloid cells 1 day after gavage of ^3^H-retinol. Cells were recovered from conventional and antibiotic-treated wild-type and *Saa*^-/-^ mice. *Saa*^-/-^ mice lack all four mouse *Saa* isoforms.^31,42^ **(D)** LC-UV detection of retinol in LP CD11c^+^ myeloid cells isolated as in (B). Representative LC-UV chromatograms are shown. **(E)** LC-UV chromatograms were used to determine relative retinol levels in LP CD11c^+^ myeloid cells. Retinol levels were quantified by LC-UV and normalized to protein. The average of the values in conventional wild-type mice was set to 1; antibiotic-treated wild-type, *Saa*^-/-^, and antibiotic-treated *Saa*^-/-^ values were expressed relative to this average. Each point represents one mouse. **(F)** Retinol dehydrogenase (RDH) and retinaldehyde dehydrogenase (RALDH) are enzymes that act sequentially to convert retinol to RA in myeloid cells. The ALDEFLUOR^TM^ assay measures RALDH activity based on the conversion of BODIPY-aminoacetaldehyde (BAAA) to fluorescent BODIPY-aminoacetate (BAA). BAA (Aldefluor) is detected in cells by flow cytometry. **(G)** Representative flow cytometry histograms of Aldefluor detection in LP CD11c^+^ myeloid cells. Diethylaminobenzaldehyde (DEAB) is a specific inhibitor of ALDH activity. DEAB-treated cells were used as a negative control. **(H)** Quantification of Aldefluor activity in (G). **(I)** Schematic of the Villin-*Saa1* transgenic mouse model (*Saa1* Tg). *Saa1* is cloned downstream of the *Vil1* (Villin) gene promoter, which drives gene expression in intestinal epithelial cells. **(J)** qPCR detection of *Saa1* mRNA in the small intestines of conventional and antibiotic-treated wild-type and *Saa1* Tg mice. **(K)** Immunofluorescence detection of SAA1/2 in the small intestines of conventional and antibiotic-treated wild-type and *Saa1* Tg mice. Scale bars = 100 μm. **(L)** Quantification of SAA1/2 immunofluorescence in (K). Mean fluorescence intensity (MFI) of SAA detection was determined for each villus across three fields per mouse. n=3 mice per group. **(M)** ³H-retinoid flux to LP CD11c^+^ myeloid cells from conventional and antibiotic-treated wild-type and *Saa1* Tg mice 1 day after gavage of ^3^H-retinol. **(N)** LC-UV detection of retinol in LP CD11c^+^ myeloid cells from conventional and antibiotic-treated wild-type and *Saa1* Tg mice. Representative LC-UV chromatograms are shown. **(O)** LC-UV chromatograms were used to determine relative retinol levels in LP CD11c^+^ myeloid cells. Retinol levels were quantified by LC-UV and normalized to protein. The average of the values in conventional wild-type mice was set to 1; antibiotic-treated wild-type, *Saa1* Tg, and antibiotic-treated *Saa1* Tg values were expressed relative to this average. Each point represents one mouse. **(P)** Representative flow cytometry histograms of Aldefluor detection in LP CD11c^+^ myeloid cells. DEAB-treated cells were used as a negative control. **(Q)** Quantification of Aldefluor activity in (P). **(R)** Summary schematic showing how the microbiota initiates retinoid delivery to intestinal myeloid cells by inducing SAA expression in epithelial cells. LP, lamina propria; cpm, counts per minute; Conv, conventional; Abx, antibiotic-treated; IEC, intestinal epithelial cell; MFI, median fluorescence intensity; LC-UV, liquid chromatography–ultraviolet spectroscopy; A.U., absorbance units; min, minutes; RDH, retinol dehydrogenase; RALDH, retinaldehyde dehydrogenase. n=5 mice per group. Data are representative of at least two independent experiments. Means ± SEM are plotted. \**p* < 0.05; \*\**p* < 0.01; \*\*\**p* < 0.001; ns, not significant by two-tailed Student’s *t*-test. See also Figure S4.

Myeloid cell retinoid metabolism was also impaired in the absence of SAA and the microbiota. The ALDEFLUOR™ assay measures aldehyde dehydrogenase (ALDH) activity through metabolism of a fluorescent substrate and, in this context, primarily reflects activity of retinaldehyde dehydrogenases (RALDHs), the ALDH family members that generate RA (Fig. 3F).^31^ Using this assay, we found that CD11c⁺ myeloid cells from both *Saa⁻^/^⁻* and antibiotic-treated mice exhibited reduced RALDH activity, suggesting that SAA and the microbiota promote retinol metabolism to RA in these cells (Fig. 3G,H). Similarly, there was decreased expression of *Aldh1a2*, *Aldh1a3*, and *Cyp26b1*—RAR target genes—in LP CD11c⁺ cells from *Saa⁻^/^⁻* mice, mirroring antibiotic-treated controls (Fig. S4F).

We next sought to test whether SAA expression is sufficient to drive retinoid flux in the absence of the microbiota. We therefore generated Villin*-Saa1* transgenic (Tg) mice that express *Saa1* under the native *Vil1* (Villin) gene promoter, which selectively directs expression in intestinal epithelial cells (Fig. 3I).^45^ This was achieved by inserting *Saa1* cDNA into the *Vil1* 3′ UTR with a preceding p2A site to ensure bicistronic expression (Fig. S4C,D). Villin*-Saa1* Tg (*Saa1* Tg) mice preserved high levels of *Saa1* expression despite antibiotic treatment (Fig. 3J–L; Fig. S4E). This model thus uncouples *Saa1* expression from the microbiota, enabling determination of their distinct contributions to intestinal retinoid flux.

In the Villin*-Saa1* Tg mice, ^3^H-retinoid flux to LP CD11c⁺ myeloid cells reached conventional levels despite the absence of the gut microbiota (Fig. 3M). Independent LC-UV measurements likewise showed that myeloid cell retinol content in antibiotic-treated Villin*-Saa1* Tg mice was indistinguishable from that of conventional controls (Fig. 3N,O). RALDH activity and expression of *Aldh1a2*, *Aldh1a3*, and *Cyp26b1* were also similar to conventional controls (Fig. 3P,Q; Fig. S4F). Thus, while the microbiota is required to induce epithelial SAA, once expressed, SAA is sufficient to mobilize retinoids from epithelial to myeloid cells and to drive RAR-dependent gene expression in myeloid cells, even in the absence of the microbiota (Fig. 3R).

### The microbiota enables the migration of retinoid-loaded CD11c^+^ myeloid cells to the mesenteric lymph nodes through SAA

We next assessed the role of the microbiota and SAA in enabling the migration of retinol-loaded myeloid cells from the intestine to the mLNs. We first enumerated CD11c⁺ myeloid cells in both compartments. Total CD11c^+^myeloid cell numbers in the small intestine were comparable across all groups (conventional wild-type, *Saa*⁻^/^⁻, and Villin-*Saa1* Tg; and antibiotic-treated wild-type and Villin-*Saa1* Tg mice). However, mLN myeloid cell numbers were reduced in conventional *Saa*⁻^/^⁻ and antibiotic-treated wild-type mice, and this defect was rescued by transgenic *Saa1* expression in antibiotic-treated mice (Fig. 4A). Changes in absolute numbers were reflected in cell frequencies (Fig. S5B,C). These results indicate that SAA expression is both necessary and sufficient to regulate myeloid cell numbers in the mLNs, independent of the microbiota.

**Figure 4:**
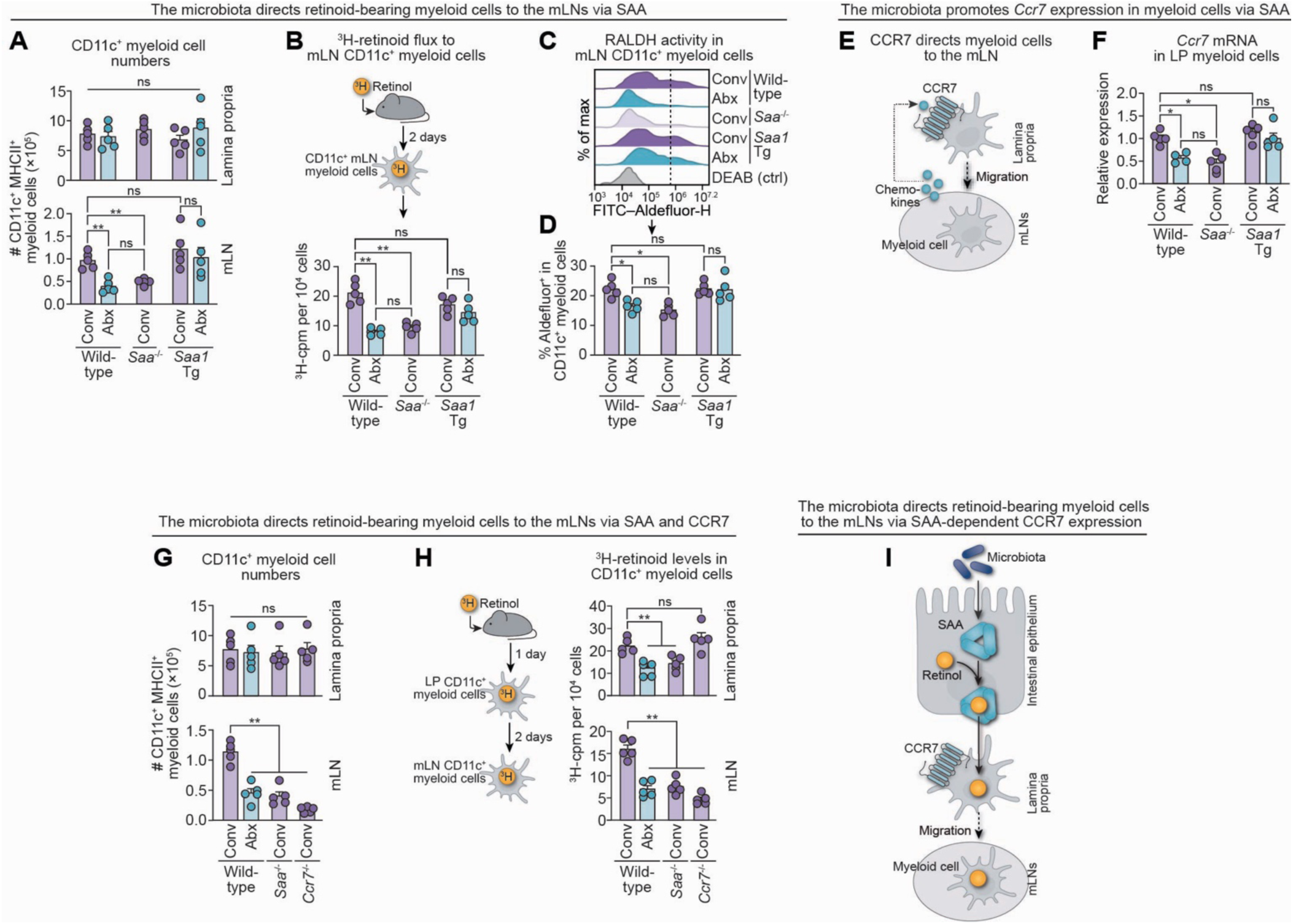
The microbiota enables the migration of retinoid-loaded CD11c^+^ myeloid cells to the mesenteric lymph nodes through SAA. **(A)** Total numbers of LP and mLN CD11c⁺ myeloid cells in conventional and antibiotic-treated wild-type, conventional *Saa^-/-^*, and conventional and antibiotic-treated *Saa1* Tg mice were determined by flow cytometry. Representative flow cytometry plots are shown in Fig. S5B. **(B)** ^3^H-retinol was administered to mice by gavage. mLN myeloid cells were isolated by magnetic sorting 2 days post-gavage, and ^3^H-cpm were quantified by scintillation counting. **(C)** Representative flow cytometry histograms of Aldefluor detection in mLN CD11c^+^ myeloid cells. DEAB-treated cells were used as a negative control. **(D)** Quantification of Aldefluor activity in (C). **(E)** Schematic of CCR7-dependent migration of CD11c⁺ myeloid cells from the LP to the mLNs. **(F)** qPCR quantification of *Ccr7* mRNA in LP CD11c⁺ myeloid cells. **(G)** Total numbers of LP and mLN CD11c⁺ myeloid cells in conventional and antibiotic-treated wild-type, conventional *Saa^-/-^*, and conventional *Ccr7^-/-^* mice were determined by flow cytometry. Representative flow cytometry plots are shown in Fig. S5B. **(H)** ³H-retinoid levels in LP CD11c⁺ myeloid cells (1 day post ^3^H-retinol gavage) and in mLN CD11c⁺ myeloid cells (2 days post ^3^H-retinol gavage). **(I)** Model showing how the microbiota directs retinoid-bearing myeloid cells to the mLNs through SAA-dependent CCR7 expression. mLNs, mesenteric lymph nodes; Conv, conventional; Abx, antibiotic-treated; cpm, counts per minute; LP, lamina propria. n=5 mice per group. Data are representative of at least two independent experiments. Means ± SEM are plotted. \**p* < 0.05; \*\**p* < 0.01; ns, not significant by two-tailed Student’s *t*-test. See also Figure S5.

We next performed ³H-retinol tracer experiments to assess retinoid flux to mLN myeloid cells. Two days after ^3^H-retinol gavage, ^3^H-retinoid levels in mLN myeloid cells were reduced in *Saa⁻^/^⁻* and antibiotic-treated wild-type mice as compared to conventional wild-type controls. However, transgenic *Saa1* expression was sufficient to restore ^3^H-retinoid levels in mLN myeloid cells to conventional levels, even in the absence of the microbiota (Fig. 4B). Correspondingly, RALDH activity in mLN myeloid cells was diminished in conventional *Saa⁻^/^⁻* and antibiotic-treated wild-type mice but restored in antibiotic-treated Villin*-Saa1* Tg mice (Fig. 4C,D). Finally, RA-dependent gene expression was reduced in the absence of SAA or the gut microbiota but was restored by *Saa1* overexpression even in the absence of the gut microbiota (Fig. S5A). These findings demonstrate that SAA is both necessary and sufficient for the migration of retinoid-loaded myeloid cells to the mLNs and that it rescues this process in the absence of the microbiota.

C-C chemokine receptor 7 (CCR7) is a G protein–coupled receptor that directs myeloid cell migration from the small intestine to the mLNs by sensing lymph node–derived chemokines (Fig. 4E).^46,47^ *Ccr7* transcripts were reduced in lamina propria myeloid cells from *Saa*⁻^/^⁻ and antibiotic-treated wild-type mice but remained high in Villin-*Saa1* Tg mice lacking a microbiota, demonstrating that SAA is both necessary and sufficient to induce myeloid cell *Ccr7* expression (Fig. 4F). Consistent with this finding, *Ccr7*⁻^/^⁻ mice harbored normal numbers of lamina propria myeloid cells but showed a marked deficiency of mLN myeloid cells, a phenotype that mirrored *Saa*⁻^/^⁻ and antibiotic-treated wild-type mice (Fig. 4G; Fig. S5B,C). In ³H-retinol tracer experiments, *Ccr7*⁻^/^⁻ myeloid cells acquired retinoids normally in the intestine but failed to deliver them to the mLNs, phenocopying *Saa*⁻^/^⁻ and antibiotic-treated wild-type mice (Fig. 4H). Importantly, our studies in Villin-*Saa1* Tg mice show that, once expressed, SAA alone is sufficient to induce *Ccr7* expression in intestinal myeloid cells, even in the absence of the microbiota. This enables the migration of these cells from the small intestine to the mLNs and the transport of retinoids between these compartments (Fig. 4I).

### Microbiota-initiated vitamin A flux promotes CD4^+^ T cell transcriptional maturation and intestinal homing

We next investigated whether the intestinal microbiota promotes vitamin A flux to developing mLN CD4⁺ T cells in an SAA-dependent manner. To test this, we measured ^3^H-retinoid levels in mLN CD4⁺ T cells three days after gavage (Fig. 5A). Both *Saa⁻^/^⁻* and antibiotic-treated wild-type mice exhibited reduced ^3^H-retinoid flux to mLN CD4⁺ T cells compared to conventional wild-type controls (Fig. 5B). Transgenic expression of *Saa1* in antibiotic-treated mice failed to restore T cell retinoid levels to those of conventional mice (Fig. 5C). Consistent with these findings, expression of RAR target genes—*Ccr9*, *Crabp2*, and *Cyp26b1*—was reduced in mLN CD4⁺ T cells recovered from *Saa⁻^/^⁻* and antibiotic-treated wild-type mice and remained persistently low in antibiotic-treated Villin*-Saa1* Tg mice (Fig. S6A). These results suggest that while SAA is necessary for retinoid flux to mLN T cells, it is not sufficient in the absence of the microbiota.

**Figure 5:**
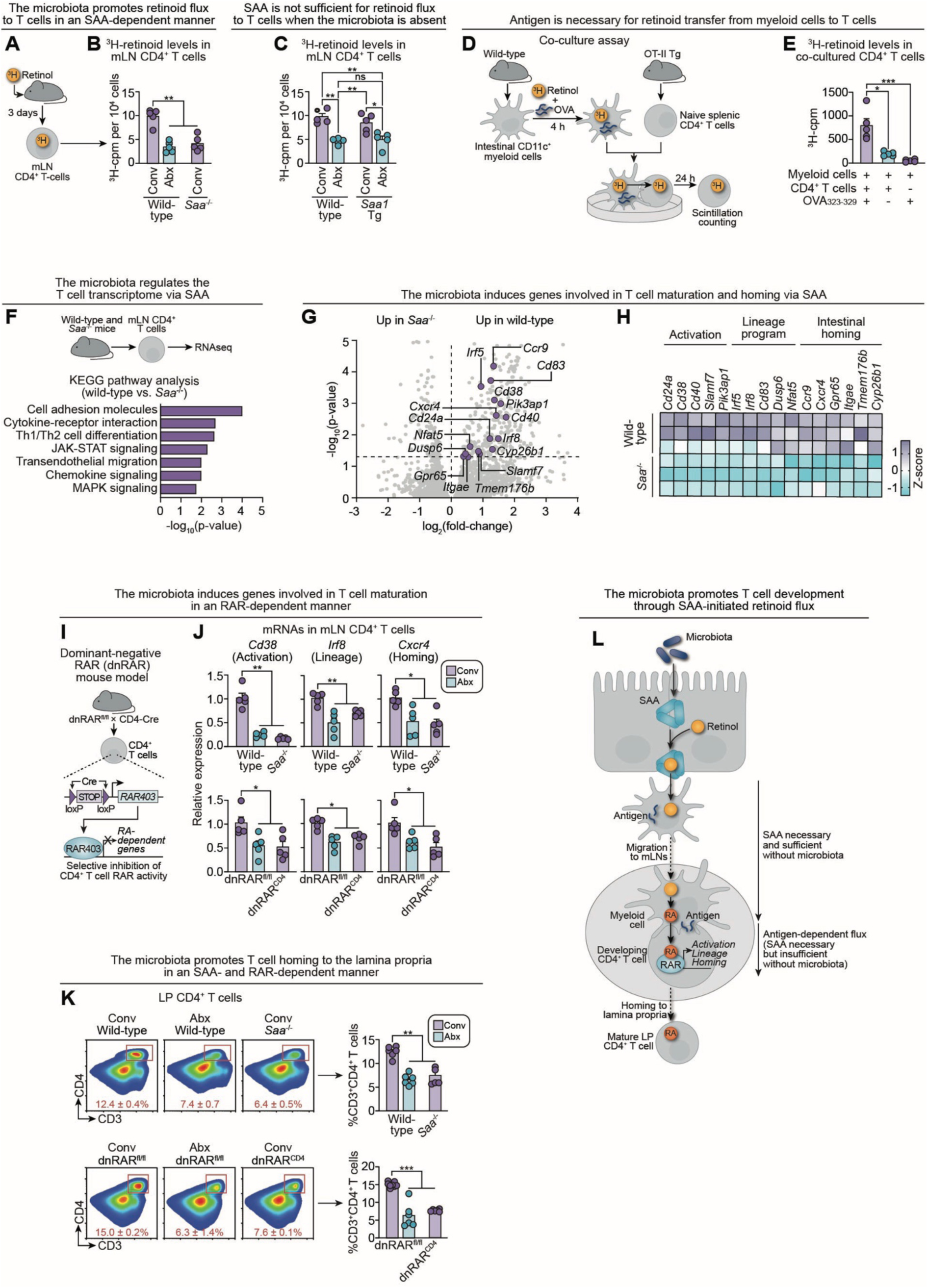
Microbiota-initiated vitamin A flux promotes CD4^+^ T cell transcriptional maturation and intestinal homing. (A) Measurement of ³H-retinoid flux to mLN CD4^+^ T cells. ^3^H-retinol was administered to mice by gavage. mLN CD4^+^ T cells were isolated by magnetic sorting 3 days post-gavage, and ^3^H-cpm were quantified by scintillation counting. (B) ³H-retinoid levels in mLN CD4⁺ T cells 3 days after gavage of ^3^H-retinol into conventional and antibiotic-treated wild-type mice and conventional *Saa*^-/-^ mice. (C) ^3^H-retinoid levels in mLN CD4^+^ T cells 3 days after gavage of ^3^H-retinol into conventional and antibiotic-treated wild-type and *Saa1* Tg mice. (D) Schematic of co-culture assay testing whether antigen is required for retinoid transfer from intestinal CD11c⁺ myeloid cells to CD4⁺ T cells. CD11c⁺ cells were pulsed with OVA_323-339_ peptide and ^3^H-retinol for 4 hours, then co-cultured with naïve splenic CD4⁺ T cells harvested from OT-II transgenic mice. Transfer of ^3^H-retinoids into the T cells was measured by scintillation counting after 24 hours. (E) ^3^H-retinoid levels in CD4⁺ T cells after 24-hour co-culture with intestinal CD11c⁺ myeloid cells with or without OVA_323-339_. ^3^H-retinoid levels in OVA-pulsed myeloid cells cultured alone for 24 hours were measured as a control. (F) RNA-seq of isolated mLN CD4⁺ T cells from wild-type and *Saa*^-/-^ mice. Kyoto Encyclopedia of Genes and Genomes (KEGG) pathway enrichment analysis of differentially expressed genes is shown. (G) Volcano plot highlighting differentially expressed genes associated with T cell activation, lineage programming, and intestinal homing. (H) Heatmap of differentially expressed genes associated with T cell activation, lineage programming, and intestinal homing. (I) Selective disruption of RAR activity in CD4^+^ T cells. Dominant-negative RAR (dnRAR) knock-in mice carry a neomycin resistance gene with three *loxP*-flanked polyadenylation sequences (STOP) located upstream of an open reading frame encoding a dominant negative (dn)RAR. The dnRAR is a mutated human RARα isoform (RAR403) that lacks the ligand-dependent transactivation domain and thus inhibits the activity of all three mouse RAR isoforms (RARα, β, and γ).^79^ Breeding to CD4-Cre mice results in selective expression of the dnRAR in CD4^+^ T cells and thus selective inhibition of RAR activity in these cells. (J) qPCR analysis of select genes related to T cell activation, lineage programming, and intestinal homing in mLN CD4⁺ T cells. (K) Representative flow cytometry plots and quantification of CD4⁺ T cell frequencies in the small intestinal lamina propria. (L) Model of how the microbiota promotes intestinal T cell development through SAA-initiated retinoid flux. mLN, mesenteric lymph nodes; LP, lamina propria; Conv, conventional; Abx, antibiotic-treated; cpm, counts per minute; OVA_323-339_, Ovalbumin peptide 323-339; RAR, retinoic acid receptor; RA, retinoic acid. n=3–8 mice per group. Data are representative of at least two independent experiments. Means ± SEM are plotted. \**p* < 0.05; \*\**p* < 0.01; \*\*\**p* < 0.001; ns, not significant by two-tailed Student’s *t*-test. See also Figure S6.

To better define the mechanistic requirements for retinoid transfer between myeloid cells and T cells, we used the OT-II system—an antigen-restricted model in which CD4⁺ T cells recognize OVA₃₂₃–₃₃₉ peptide presented by myeloid cells in *ex vivo* co-culture.^48,49^ Naïve splenic OT-II CD4⁺ T cells were co-cultured with intestinal myeloid cells preloaded with ³H-retinol and either pulsed or not pulsed with OVA₃₂₃–₃₃₉ (Fig. 5D). T cells co-cultured with OVA peptide-pulsed myeloid cells exhibited greater ³H-retinoid uptake, indicating that the presence of antigen enhances retinoid transfer (Fig. 5E). While OVA is not a microbial antigen, its ability to facilitate transfer in this system, together with the lack of observable transfer in antibiotic-treated mice, suggests that microbial antigen may provide the physiological trigger for myeloid-to-T cell retinoid exchange. This framework could explain why, in the absence of the microbiota, transgenic *Saa1* expression alone is insufficient to drive retinoid delivery to T cells in Villin-*Saa1* Tg mice. Importantly, intestinal myeloid cells from wild-type, *Saa⁻^/^⁻*, and Villin*-Saa1* Tg mice displayed similar co-stimulatory molecule expression and equivalent capacity to activate OT-II CD4⁺ T cells *in vitro* following OVA₃₂₃–₃₃₉ pulsing (Fig. S6B–E). In contrast, intestinal myeloid cells from antibiotic-treated wild-type and Villin*-Saa1* Tg mice exhibited impairments in co-stimulatory molecule expression and OT-II CD4⁺ T cell activation. These findings indicate that SAA is dispensable for myeloid cell antigen presentation, whereas microbial exposure is necessary. Taken together, these data suggest that SAA and microbial antigen drive distinct steps of retinoid flux through the myeloid–T cell axis.

To assess the functional consequences of impaired intestinal vitamin A flux to mLN CD4⁺ T cells, we performed RNA sequencing on mLN CD4⁺ T cells from wild-type and *Saa⁻^/^⁻* mice (Fig. 5F). By preserving microbial exposure while disrupting vitamin A flux, this comparison isolates the specific contribution of retinoid availability to T cell development. Kyoto Encyclopedia of Genes and Genomes (KEGG) pathway analysis of differentially expressed genes revealed enrichment of transcriptional programs governing T cell activation (JAK-STAT signaling, MAPK signaling)^50,51^, lineage specification (cytokine–cytokine receptor interaction, Th1/Th2 differentiation)^52,53^, and intestinal homing (chemokine signaling, leukocyte transendothelial migration, cell adhesion molecules)^19,54^, suggesting defects in intestinal localization and functional maturation of CD4⁺ T cells in the absence of SAA (Fig. 5F). Closer analysis revealed specific genes within these pathways that were downregulated in *Saa⁻^/^⁻* mice (Fig. 5G,H).

To validate these transcriptional changes, we quantified representative transcripts across functional categories by qPCR of mLN CD4⁺ T cells from wild-type, *Saa⁻^/^⁻*, and antibiotic-treated wild-type mice. Expression of activation, lineage-associated, and intestinal homing genes was reduced in both SAA-deficient and microbiota-depleted mice, indicating that the microbiota regulates CD4⁺ T cell maturation through intestinal SAA (Fig. 5J).

To test the role of RAR signaling in regulating this microbiota-driven transcriptional maturation, we employed a dominant-negative retinoic acid receptor (dnRAR) mouse model crossed to CD4-Cre transgenic mice, selectively ablating RAR activity in CD4⁺ T cells (Fig. 5I).^20,55,56^ CD4⁺ T cells from dnRAR^CD4^ mice expressed the dnRAR transgene and exhibited reduced expression of canonical RAR target genes, confirming effective blockade of RAR signaling (Fig. S6F,G). Strikingly, gene expression defects observed in *Saa⁻^/^⁻* mice were recapitulated in dnRAR^CD4^ mice, supporting a T cell-intrinsic requirement for RAR signaling in mediating these transcriptional programs (Fig. 5J). Antibiotic-treated dnRAR^fl/fl^ mice displayed similar defects, suggesting that the microbiota promotes RAR-dependent CD4⁺ T cell maturation and intestinal localization. Flow cytometry revealed a downstream functional consequence of these transcriptional changes: reduced accumulation of CD4⁺ T cells in the small intestines of antibiotic-treated wild-type, *Saa⁻^/^⁻*, and dnRAR^CD4^ mice compared to conventional wild-type mice (Fig. 5K).

Together, these data show that SAA-dependent retinoid flux activates T cell–intrinsic RAR signaling, which in turn drives transcriptional programs required for CD4⁺ T cell functional maturation and intestinal homing. Our findings from the Villin-*Saa1* Tg mice further delineate two distinct segments of the retinoid flux pathway: an SAA-dependent phase that is microbiota-independent and a subsequent phase that requires both SAA and microbial antigen (Fig. 5L).

### The microbiota activates a myeloid cell–ILC3–epithelial cell signaling relay to initiate vitamin A flux

The microbiota induces expression of many IEC genes—including *Saa* genes—via a multicellular relay triggered by microbe-associated molecular patterns (MAMPs), such as lipopolysaccharide and flagellin^3,57–59.^ MAMPs engage Toll-like receptors (TLRs) on CD11c^+^ myeloid cells, signaling through the adaptor MYD88 to drive interleukin-23 (IL-23) production. IL-23 activates group 3 innate lymphoid cells (ILC3), which secrete IL-22 to stimulate IEC STAT3 and thereby induce transcription of targets that include *Saa* genes.^3,57–59^

To test whether this circuit controls microbiota-induced retinoid flux, we first examined the requirement for myeloid cell *Myd88*. Deleting *Myd88* in CD11c⁺ myeloid cells (*Myd88*^ΔCD11c^), but not in IECs (*Myd88*^ΔIEC^), reduced intestinal *Saa1* expression (Fig. 6A). Consistent with this reduction, LP myeloid cell retinol content and ^3^H-retinoid flux to mLN T cells were both diminished in *Myd88*^ΔCD11c^ mice (Fig. 6B,C). These defects phenocopied antibiotic-treated *Myd88^fl/fl^* controls (Fig. 6A–C), indicating that myeloid cell *Myd88* is required for microbiota-driven retinoid flux.

**Figure 6:**
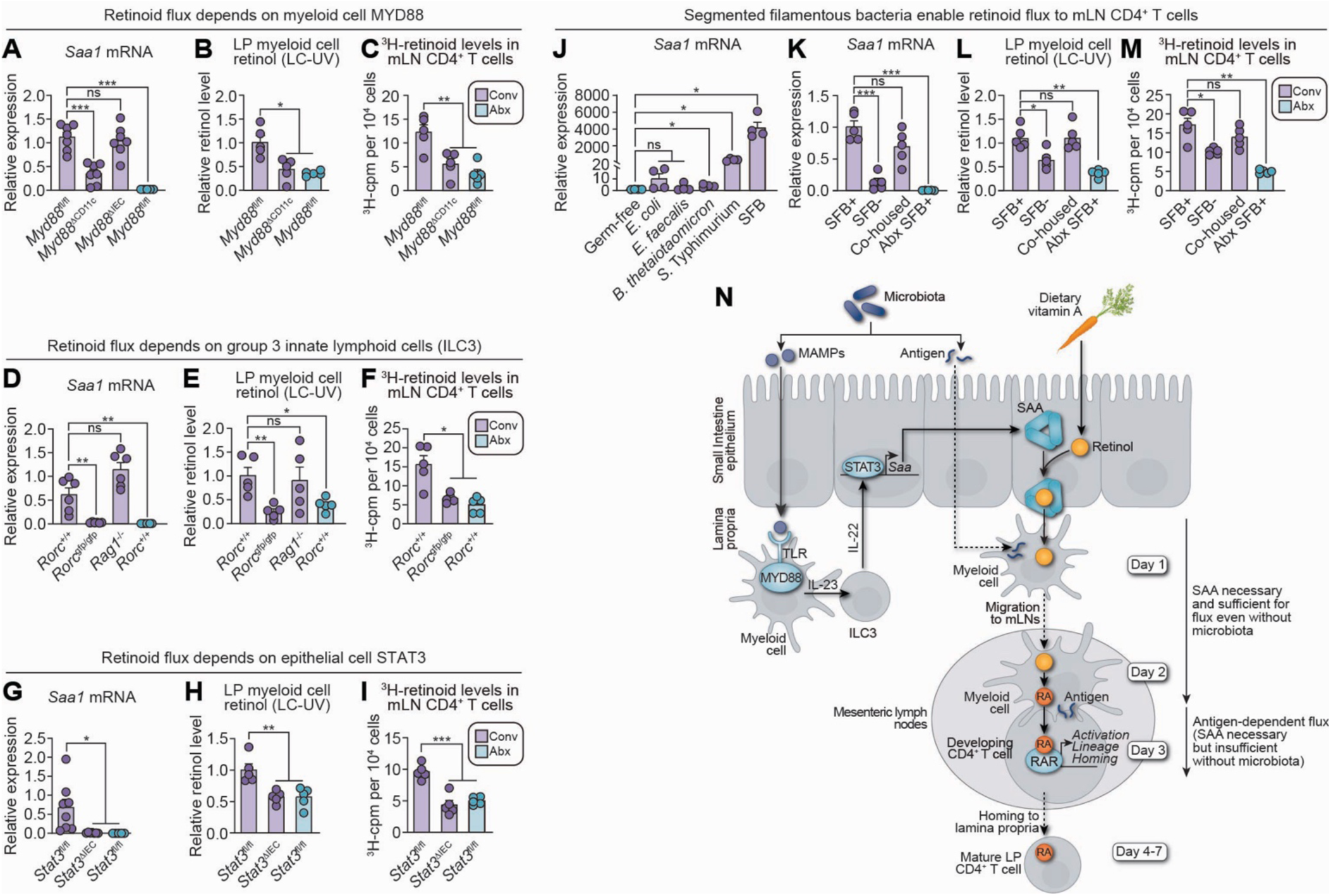
The microbiota activates a myeloid cell–ILC3–epithelial cell signaling relay to initiate vitamin A flux. **(A-C)** The role of the Toll-like receptor signaling adaptor MYD88 in microbiota-driven retinoid flux was assessed by analyzing conventional *Myd88*^fl/fl^, *Myd88*^ΔCd11c^ (myeloid cell deletion), and *Myd88*^ΔIEC^ (IEC deletion) mice, as well as antibiotic-treated *Myd88*^fl/fl^ mice. **(D-F)** The role of group 3 innate lymphoid cells (ILC3) in microbiota-driven retinoid flux was assessed by analyzing conventional *Rorc*^+/+^, *Rorc*^gfp/gfp^ (lacking ILC3s and Th17 CD4^+^ T cells), and *Rag1*^-/-^ mice (lacking all T cells), as well as antibiotic-treated *Rorc*^+/+^ mice. **(G-I)** The role of epithelial cell STAT3 in microbiota-driven retinoid flux was assessed by analyzing conventional *Stat3*^fl/fl^ and *Stat3*^ΔIEC^ mice (with an IEC-specific deletion of *Stat3*), and antibiotic-treated *Stat3*^fl/fl^ mice. **(A,D,G)** qPCR quantification of small intestine *Saa1* mRNA in the mouse groups listed above. **(B,E,H)** Retinol levels were measured by LC-UV analysis of LP CD11c⁺ myeloid cells from the mouse groups listed above. **(C,F,I)** ³H-retinoid flux to mLN CD4⁺ T cells 3 days after ^3^H-retinol gavage of the mouse groups listed above. **(J)** qPCR quantification of small intestine *Saa1* mRNA in germ-free mice and germ-free mice monocolonized for 3 days with *Escherichia coli*, *Enterococcus faecalis*, *Bacteroides thetaiotaomicron*, *Salmonella typhimurium*, or segmented filamentous bacteria (SFB). **(K–M)** The impact of SFB colonization on microbiota-driven retinoid flux was assessed by analyzing SFB^+^ conventional mice (SFB^+^), SFB⁻ conventional mice (SFB^-^), co-housed mice (SFB^-^ mice cohoused with SFB^+^ mice for 14 days), and antibiotic-treated SFB^+^ mice (Abx SFB^+^). **(K)** qPCR quantification of small intestine *Saa1* mRNA in the mouse groups listed above. **(L)** Retinol levels were measured by LC-UV analysis of LP CD11c⁺ myeloid cells from the mouse groups listed above. **(M)** ³H-retinoid flux to mLN CD4⁺ T cells 3 days after ^3^H-retinol gavage of the mouse groups listed above. **(N)** Model of how the microbiota directs retinoid flux to mLN T cells, with distinct roles for microbe-associated molecular patterns (MAMPs) versus antigen. First, microbiota MAMPs trigger TLR-MYD88 signaling on lamina propria myeloid cells. This signal activates ILC3s and culminates in STAT3-mediated activation of *Saa* gene expression in IECs. SAAs transfer retinol from IECs to lamina propria myeloid cells, and the retinol-loaded myeloid cells migrate to the mesenteric lymph nodes. Myeloid cells convert retinol to retinoic acid, which is transferred to developing CD4^+^ T cells, where it activates a transcriptional program that promotes T cell maturation and migration to the intestinal lamina propria. Expression of SAA in the absence of the microbiota is sufficient to drive retinol loading of lamina propria myeloid cells and their migration to the mLNs. However, antigen is required for the transfer of retinoic acid from myeloid cells to T cells, where it engages RARs to activate a developmental program. Through this pathway, the microbiota mobilizes a developmental signal across tissue compartments to program intestinal CD4^+^ T cell maturation and homing. LP, lamina propria; mLN, mesenteric lymph nodes; LC–UV, liquid chromatography–ultraviolet spectroscopy; MAMPs, microbe-associated molecular patterns; RA, retinoic acid; RAR, retinoic acid receptor. n=5–8 mice per group. All data are representative of at least two independent experiments. Means ± SEM are plotted. \**p* < 0.05; \*\**p* < 0.01; \*\*\**p* < 0.001; ns, not significant as determined by two-tailed Student’s *t*-test.

We next assessed ILC3 involvement using *Rorc^gfp/gfp^* mice,^60^ which lack RORγt⁺ cells, including ILC3s and Th17 cells. Relative to *Rorc*^+/+^ controls, *Rorc^gfp/gfp^* mice displayed reduced IEC *Saa1* expression, lower myeloid cell retinol, and impaired ^3^H-retinoid flux to mLN T cells (Fig. 6D–F). By contrast, *Rag1⁻^/^⁻* mice,^61^ which lack T cells but retain ILC3s, maintained normal *Saa1* expression and normal myeloid cell retinol levels, implicating ILC3s rather than Th17s in initiating retinoid flux (Fig. 6D,E). Antibiotic-treated *Rorc^+/+^* controls showed flux defects similar to those of *Rorc^gfp/gfp^* mice (Fig. 6F), supporting an essential role for ILC3s in microbiota-initiated retinoid flux.

Finally, we examined epithelial STAT3. Mice with an IEC-specific deletion of *Stat3* (*Stat3^ΔIEC^*) exhibited decreased *Saa1* expression, reduced myeloid cell retinol, and impaired ^3^H-retinoid flux to mLN T cells, mirroring antibiotic-treated *Stat3*^fl/fl^ controls (Fig. 6G–I). Thus, epithelial *Stat3* is required for microbiota-initiated retinoid flux. Together, these results indicate that the microbiota engages a myeloid–ILC3–epithelial circuit to promote epithelial SAA expression and initiate retinoid flux to mLN T cells (Fig. 6N).

To assess whether certain bacterial species selectively drive retinoid flux, we monocolonized germ-free mice with five intestinal bacterial species. Segmented filamentous bacteria (SFB)—members of the intestinal microbiota of rodents, non-human primates, and humans—robustly induced epithelial *Saa1* expression, consistent with prior work (Fig. 6J).^34,62–66^ In conventional SFB-positive (SFB^+^) mice, *Saa1* expression, myeloid cell retinol content, and ^3^H-retinoid flux to mLN T cells were elevated relative to antibiotic-treated SFB^+^ mice and conventional SFB⁻ controls (Fig. 6K–M). Introducing SFB into SFB^-^ mice by co-housing rescued *Saa1* expression, myeloid cell retinol, and ^3^H-retinoid flux to mLN T cells (Fig. 6K–M). Notably, SFB⁻ mice still exhibited greater retinoid flux than antibiotic-treated mice, indicating that SFB amplifies—but does not fully account for—microbiota-induced retinoid flux. These findings identify SFB as a potent enhancer, but not the sole driver, of intestinal retinoid flux among commensal bacteria.

### Microbiota-initiated vitamin A flux supports maturation of the intestinal immune system during postnatal development

To define the physiological context in which microbiota-driven vitamin A flux is engaged, we examined early postnatal development—a period marked by dietary transition from maternal milk to solid food and a concurrent surge in gut microbial colonization.^67–69^ A marked expansion of the gut microbiota occurs during this developmental window, in parallel with an expansion in intestinal immune defenses that maintain mucosal homeostasis.^67–72^ We hypothesized that microbiota-induced SAA expression and retinoid flux might be activated during this window to support maturation of the intestinal adaptive immune system.

We first assessed intestinal SAA gene expression in conventional mice at 2, 4, and 6 weeks of age. qPCR of small intestinal tissue revealed a progressive increase in *Saa1* expression during early postnatal development, while expression of genes encoding other retinol-binding proteins (*Rbp1*, *Rbp4*) remained unchanged or declined (Fig. 7A; Fig. S7A). To determine if this increase corresponds with retinol transfer to lamina propria myeloid cells, we measured retinol levels in LP myeloid cells from conventional wild-type, conventional *Saa⁻^/^⁻*, and germ-free wild-type mice across the same timepoints. LC-UV analysis showed a marked increase in retinol content in LP myeloid cells from conventional wild-type mice during postnatal development (Fig. 7B,C). This increase was blunted in both *Saa⁻^/^⁻* and germ-free wild-type mice, implicating both the microbiota and SAA in this process. Notably, IEC retinol content did not change with age (Fig. S7B), indicating that increased myeloid cell retinol is not due to enhanced absorption.

**Figure 7:**
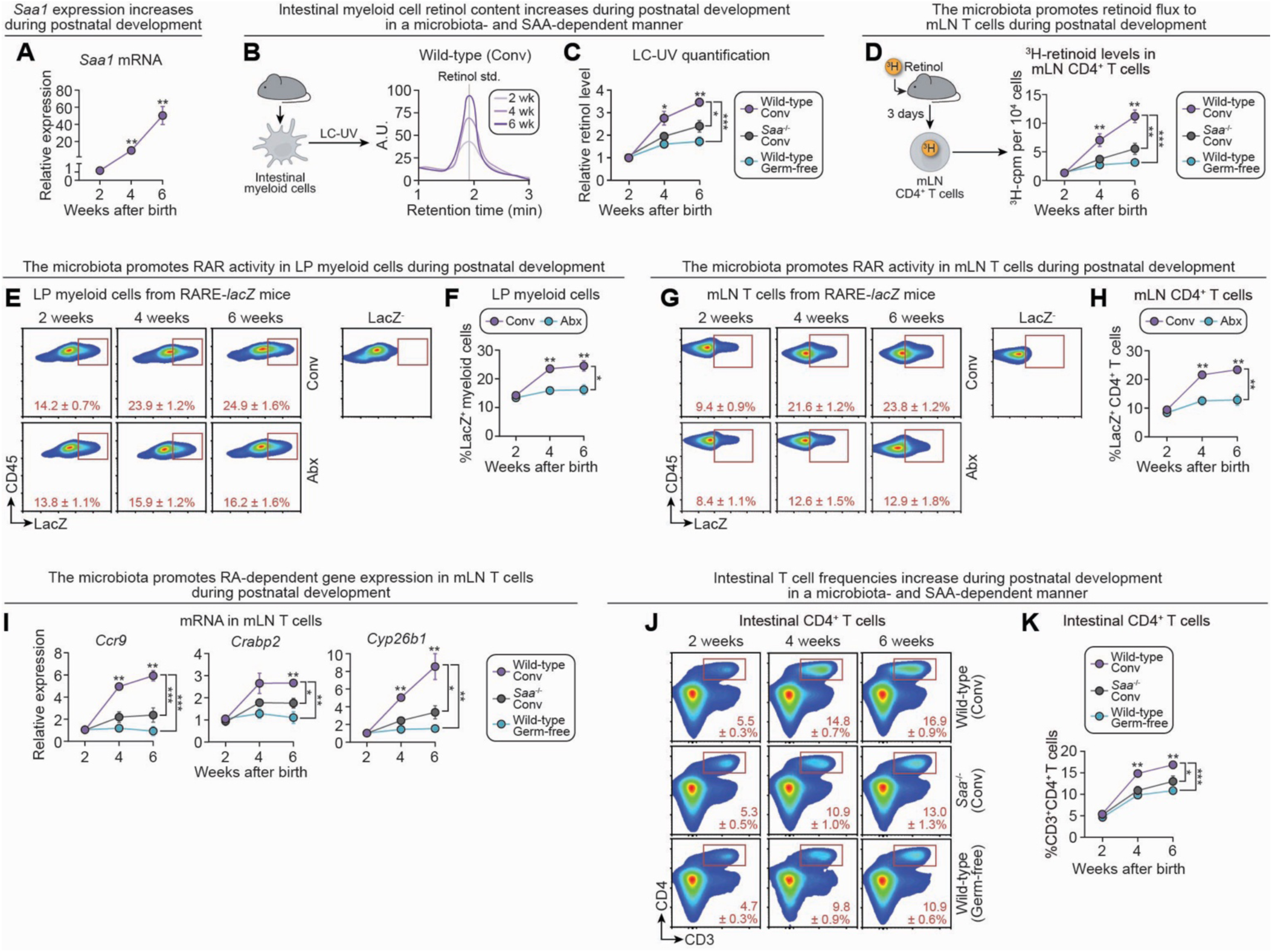
Microbiota-initiated vitamin A flux supports maturation of the intestinal immune system during postnatal development. **(A)** qPCR analysis of *Saa1* mRNA in the small intestines of 2-, 4-, and 6-week-old conventional wild-type mice. **(B)** LC-UV detection of retinol in lamina propria CD11c^+^ myeloid cells isolated from conventional wild-type, conventional *Saa^-^*^/-^, and germ-free wild-type mice at 2, 4, and 6 weeks. Representative chromatograms are shown. **(C)** LC-UV chromatograms were used to determine relative retinol levels in LP CD11c^+^ myeloid cells. Retinol levels were quantified by LC-UV and normalized to protein. The average of the values in conventional wild-type mice was set to 1; conventional *Saa*^-/-^ and germ-free wild-type values were expressed relative to this average. **(D)** ³H-retinoid flux to mLN CD4⁺ T cells 3 days after ^3^H-retinol gavage. **(E)** Representative flow cytometry plots showing detection of LacZ in lamina propria CD11c^+^ myeloid cells in conventional and antibiotic-treated RARE-*lacZ* mice at 2, 4, and 6 weeks. Myeloid cells were gated as previously described (live CD45⁺CD3⁻B220⁻CD11c⁺MHCII⁺), and LacZ activity was then plotted relative to CD45 expression. **(F)** Quantification of LacZ^+^ cells from (E). **(G)** Representative flow cytometry plots showing detection of LacZ in mLN CD4^+^ T cells in conventional and antibiotic-treated RARE-*lacZ* mice at 2, 4, and 6 weeks. CD4^+^ T cells were gated as previously described (live CD45⁺B220⁻CD3⁺CD4⁺), and LacZ activity was then plotted relative to CD45 expression. **(H)** Quantification of LacZ^+^ cells from (G). **(I)** qPCR analysis of RA-dependent genes (*Ccr9*, *Crabp2*, *Cyp26b1*) in mLN CD4⁺ T cells at 2, 4, and 6 weeks in conventional wild-type, conventional *Saa^-/-^*, and germ-free wild-type mice. **(J)** Representative flow cytometry plots of CD4^+^ T cells from the small intestines of conventional wild-type, conventional *Saa^-/-^*, and germ-free wild-type mice at 2, 4, and 6 weeks. **(K)** Quantification of the frequency of intestinal CD4⁺ T cells in conventional wild-type, conventional *Saa^-/-^*, and germ-free wild-type mice at each time point. In panels A, C, D, F, H, I, and K, asterisks above data points indicate significance relative to the 2-week time point, and asterisks on the right margin indicate significance between mouse groups at the 6-week time point. LC-UV, liquid chromatography–ultraviolet spectroscopy; A.U., absorbance units; min, minutes; Conv, conventional; Abx, antibiotic-treated; LP, lamina propria; mLN, mesenteric lymph nodes; RA, retinoic acid; RAR, retinoic acid receptor. n=5 mice per group per time point. Data are representative of at least two independent experiments. Means ± SEM are plotted. \**p* < 0.05; \*\**p* < 0.01; \*\*\**p* < 0.001 by two-tailed Student’s *t*-test. See also Figure S7.

To assess retinoid flux, we administered ^3^H-retinol at 2, 4, and 6 weeks of age and measured ^3^H-retinoid levels in mLN CD4⁺ T cells three days later. mLN T cell retinoid levels progressively increased in conventional wild-type mice across the early postnatal period but remained low in both conventional *Saa⁻^/^⁻* and germ-free wild-type mice (Fig. 7D), confirming that both SAA and the microbiota are required for retinoid delivery to T cells during this period. In contrast, ^3^H-retinoid uptake by IECs remained stable across groups, further excluding differences in retinoid absorption as a confounding factor (Fig. S7C).

We next assessed RAR signaling using RARE-*lacZ* reporter mice. RAR activity in LP CD11c⁺ myeloid cells and mLN CD4⁺ T cells progressively increased during early postnatal development in conventional mice but remained low in antibiotic-treated counterparts, mirroring ^3^H-retinoid uptake patterns (Fig. 7E–H). Consistent with these results, expression of RAR target genes in mLN CD4^+^ T cells—*Ccr9*, *Crabp2*, and *Cyp26b1*—also increased during early postnatal development in conventional wild-type mice but remained low in both conventional *Saa⁻^/^⁻* and germ-free wild-type mice (Fig. 7I). These findings indicate that myeloid and T cell RAR activity rise during postnatal development and depend on both the microbiota and intestinal SAA.

Finally, flow cytometric analysis of the small intestinal lamina propria revealed a striking increase in LP CD4⁺ T cell infiltration during the early postnatal period in conventional wild-type mice, as other groups have previously observed.^67,68^ This accumulation was impaired in the absence of SAA or the microbiota (Fig. 7J,K).

Together, these findings reveal that microbiota-induced, SAA-mediated vitamin A flux is initiated during postnatal development to support CD4⁺ T cell maturation and intestinal localization. This mechanism thus promotes maturation of the adaptive immune system during a critical developmental window that encompasses the expansion of the gut microbiota and heightened barrier vulnerability.

## Discussion

The small intestine is the body’s primary site of nutrient absorption and a habitat for a dense and diverse microbial community. While the microbiota can compromise barrier integrity, it also provides essential cues that promote immune maturation.^4,5,7,10,32,33,73^ In parallel, nutrient-derived metabolites such as retinoic acid are required for T cell maturation.^19^ Yet how microbial and dietary signals converge to orchestrate adaptive immune development has previously been unclear.

Here, we have defined a microbiota-dependent pathway that routes vitamin A to the immune system through sequential transfer from epithelial cells to myeloid cells to developing CD4⁺ T cells. This microbiota-driven vitamin A flux activates RAR signaling in mLN CD4⁺ T cells and initiates transcriptional programs associated with activation, lineage specification, and mucosal homing. Retinoid transfer along the epithelial–myeloid–T cell axis increases during early postnatal development, suggesting that microbial and dietary transitions at this stage activate the retinoid flux pathway to promote intestinal T cell maturation and localization.^67,68^

A central insight from our study is the identification of distinct roles for microbiota-induced SAA and microbial antigen in delivering the vitamin A–derived developmental signal to developing intestinal CD4⁺ T cells. Although prior studies established that the microbiota induces intestinal SAA expression and that SAA binds retinol,^31,59^ how microbial antigen and SAA coordinate to deliver a nutrient-derived developmental signal to the immune system remained unclear. We show that microbiota-induced SAA mobilizes retinoids from epithelial to myeloid cells and promotes the migration of retinoid-loaded myeloid cells to mLNs independent of microbiota colonization, suggesting that this segment of the pathway does not require ongoing microbial antigen stimulation. However, SAA alone, in the absence of the microbiota, is insufficient to mediate retinoid transfer to T cells, and microbial antigen is required for this final step.

These findings reveal that the microbiota coordinates micronutrient delivery across immune compartments through distinct, stage-specific signals. The first phase of retinoid flux is driven by MAMPs through an innate immune signaling relay involving myeloid cells, ILC3s, and the IEC transcription factor STAT3. This phase results in retinol loading of lamina propria myeloid cells and their migration to the mLNs. The second phase of retinoid flux, in which retinoic acid is transferred from mLN myeloid cells to developing CD4^+^ T cells, requires antigen. Together, our results expand the current understanding of how the microbiota integrates dietary and antigenic signals to shape immune development.

SAA-mediated retinoid flux is engaged during the early postnatal period, coinciding with increased intestinal T cell infiltration characteristic of this developmental window. Early postnatal development is marked by profound environmental, dietary, and microbial changes. Around days 17–21, mice transition from maternal milk to chow in the process of weaning, introducing a new consortium of microbes to the gut.^67–69^ This expanded microbiota imposes a major antigenic challenge, requiring concurrent expansion of the immune system. In this context, the mechanism we describe becomes particularly relevant: postnatal induction of SAA expression drives retinoid flux to developing T cells, licensing their localization to the small intestine. This process ensures that the adaptive immune system matures in step with microbial exposure and can effectively protect the host from infection. Several important questions remain regarding microbiota-driven intestinal retinoid flux. First, how intestinal epithelial cells partition absorbed retinoids between systemic circulation and local immune delivery remains unknown. Although the intestine is the primary site of vitamin A uptake,^22^ only a small fraction is acquired by immune cells. Our tracer studies show that ∼2% of absorbed dietary retinoids are found in intestinal myeloid cells 24 hours post-gavage, with the vast majority entering systemic circulation. This suggests that IECs selectively route retinoids to the immune compartment. However, the mechanisms governing this allocation—and how they adapt during enteric infection or inflammation—remain to be defined.

Second, it remains unclear whether the same intestinal myeloid cells that sense microbial signals also serve as the primary recipients of epithelial retinoids. Our data show that deletion of *Myd88* in CD11c⁺ myeloid cells markedly impairs both *Saa1* induction and downstream retinoid flux, indicating that this population plays a central role in initiating the epithelial–myeloid signaling axis. At the same time, CD11c⁺ myeloid cells appear to be key recipients of epithelial retinoids, suggesting a dual function in detecting microbial cues and acquiring vitamin A metabolites. Although the CD11c⁺ compartment is heterogeneous,^74^ it would be logical for the same cells that acquire microbial antigens and MAMPs to also signal a “request” for retinoids. In this model, microbial sensing by myeloid cells triggers an IL-23–IL-22–STAT3 relay that induces epithelial SAA expression. SAA then delivers retinol back to these same myeloid cells, licensing their migration to mesenteric lymph nodes. This ensures that antigen-bearing cells are equipped with retinoids that may facilitate CCR7-dependent trafficking. While we have not yet determined whether *Ccr7* expression is RAR-dependent, this presents a compelling hypothesis for future study. Alternatively, distinct myeloid subsets may partition these functions, with some cells specialized for IL-23 production and others optimized for retinol uptake. Our previous studies showed that multiple CD11c⁺ subsets express the SAA receptor low-density lipoprotein receptor–related protein 1 (LRP1), supporting the possibility of division of labor within the compartment.^31^ Dissecting which subsets integrate microbial sensing with retinoid acquisition will be critical for understanding how the intestinal immune system coordinates antigen presentation and nutrient-derived signals during T cell priming.

Third, the mechanism by which retinoids are transferred from myeloid cells to T cells is not yet understood. Prior studies have shown that SAA can bind LRP1 on myeloid cells to facilitate retinoid uptake from epithelial cells,^31^ but whether a comparable receptor-mediated process exists between myeloid cells and T cells is unknown. Our *in vivo* data show that orally delivered ³H-retinol accumulates in myeloid cells before appearing in T cells, suggesting a directional transfer. Furthermore, in *ex vivo* co-cultures, we found that retinoids move from myeloid cells to OT-II T cells more efficiently in the presence of cognate antigen (OVA_323-339_), implying that antigen recognition enhances retinoid transfer. While OVA is not a microbial antigen, its ability to promote transfer in this reductionist system, together with the lack of transfer in antibiotic-treated mice, suggests that microbial antigen may be required. Although other microbial components such as metabolites could contribute, our data most strongly point to microbial antigen as the factor licensing retinoid movement from myeloid cells to T cells.

One possible mechanism that could explain how myeloid–T cell retinoid transfer occurs is that the immunological synapse, formed during antigen presentation, serves as a site of retinoid exchange. T cells are known to acquire plasma membrane fragments and associated proteins from antigen-presenting cells through trogocytosis,^75,76^ and retinoid transfer could occur by a similar mechanism. Alternatively, retinoid delivery may function not merely as cargo transfer but as a licensing signal—stabilizing or reinforcing the immunological synapse to permit sustained signaling and T cell differentiation. Because retinoids are hydrophobic,^77^ their diffusion between cells may be favored when membranes remain in tight,^78^ prolonged contact, as occurs at synapses formed by high-affinity T-cell receptor–MHC interactions. In this model, RA could function as a “test” molecule, linking strong antigen recognition to downstream T cell maturation. Determining whether RA movement occurs through direct contact or alternative routes remains an important direction for future study.

In summary, we identify a microbiota-dependent mechanism for vitamin A trafficking to the gut immune system, initiated by epithelial SAA and culminating in the maturation and intestinal localization of CD4⁺ T cells during early postnatal development. By uncoupling the distinct functions of the microbiota and vitamin A transport, our findings reveal how the host integrates distinct microbial and dietary cues to coordinate immune development with microbial exposure in early life.

---

**Figure S1:**
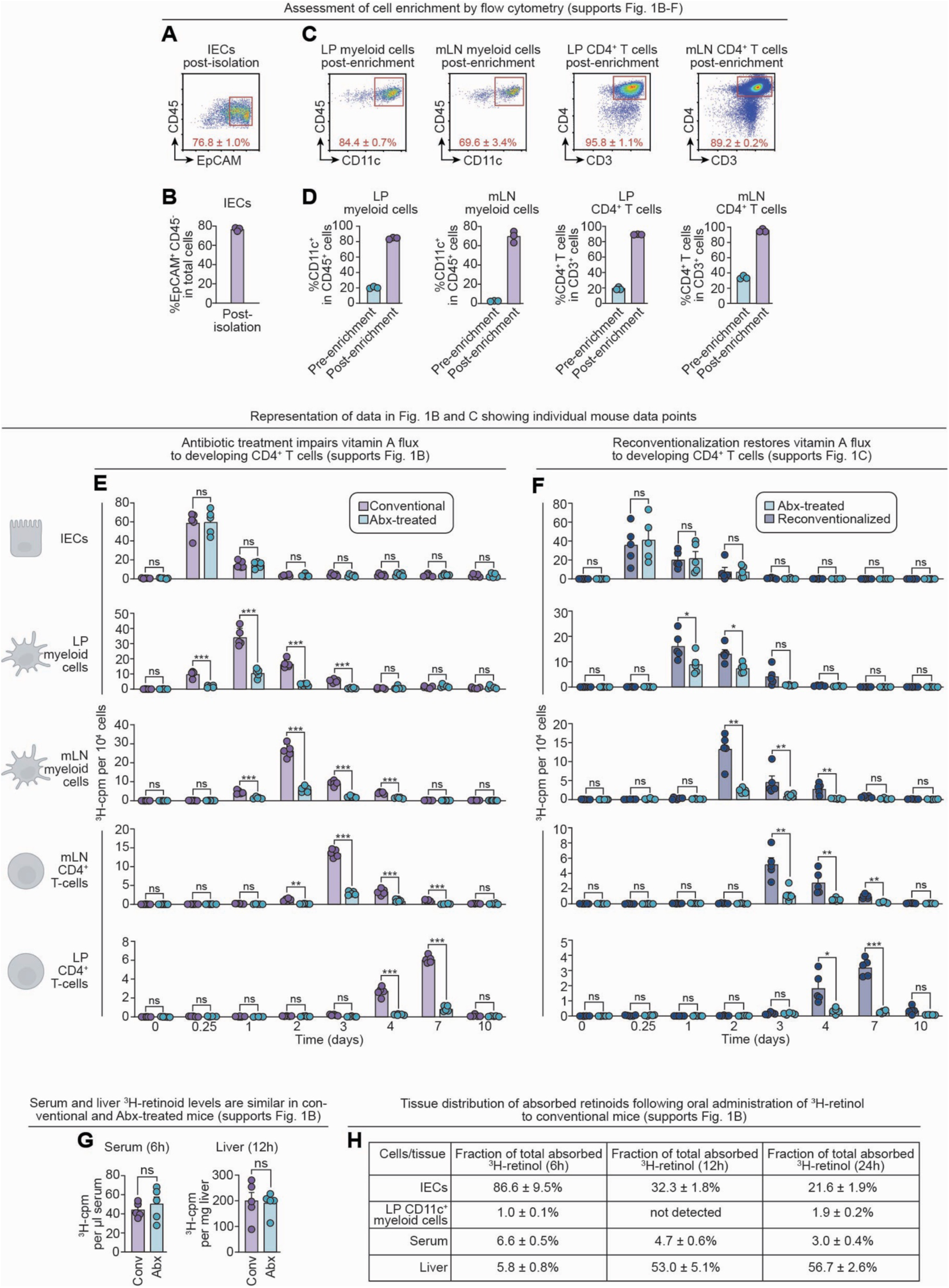
Purity of enriched immune populations, systemic ³H-retinoid uptake, and full kinetics of retinoid trafficking across the IEC–myeloid–T cell axis (supports Figure 1). **(A)** Intestinal epithelial cells (IECs) were isolated by EDTA treatment of mouse small intestinal tissue.^80^ Representative flow cytometry plots following EDTA-based isolation are shown. EpCAM marks IECs. **(B)** Quantification of IEC purity following EDTA-based isolation. **(C)** Lamina propria (LP) and mesenteric lymph node (mLN) CD11c⁺ myeloid cells and CD4⁺ T cells were isolated using magnetic bead enrichment. Representative flow cytometry plots of enriched populations are shown. **(D)** Quantification of post-enrichment purity. **(E)** Measurement of ³H-retinoid flux through IECs, LP CD11c⁺ myeloid cells, mLN CD11c⁺ myeloid cells, mLN CD4⁺ T cells, and LP CD4⁺ T cells from conventional and antibiotic-treated mice. **(F)** Measurement of ³H-retinoid flux in the same cell populations recovered from antibiotic-treated mice and mice that were reconventionalized after antibiotic treatment. **(G)** Following gavage of ³H-retinol, serum and liver samples were collected at 6h and 12h, respectively, from conventional (Conv) and antibiotic-treated (Abx) mice. ³H counts per minute (cpm) were normalized to serum volume or liver tissue mass. **(H)** Distribution of absorbed ³H-retinol among IECs, LP CD11c⁺ myeloid cells, serum, and liver at 6, 12, and 24 hours post-gavage. Values are calculated as the percentage of total ³H-retinoid content at the 6-hour time point. Values are presented as means ± SEM Conv, conventional; Abx, antibiotic-treated; ReConv, reconventionalized; cpm, counts per minute. n=3-5 mice per group. All data are representative of at least two independent experiments. Means ± SEM are plotted. \**p* < 0.05; \*\**p* < 0.01; \*\*\**p* < 0.001; ns, not significant by two-tailed Student’s *t*-test.

**Figure S2.**
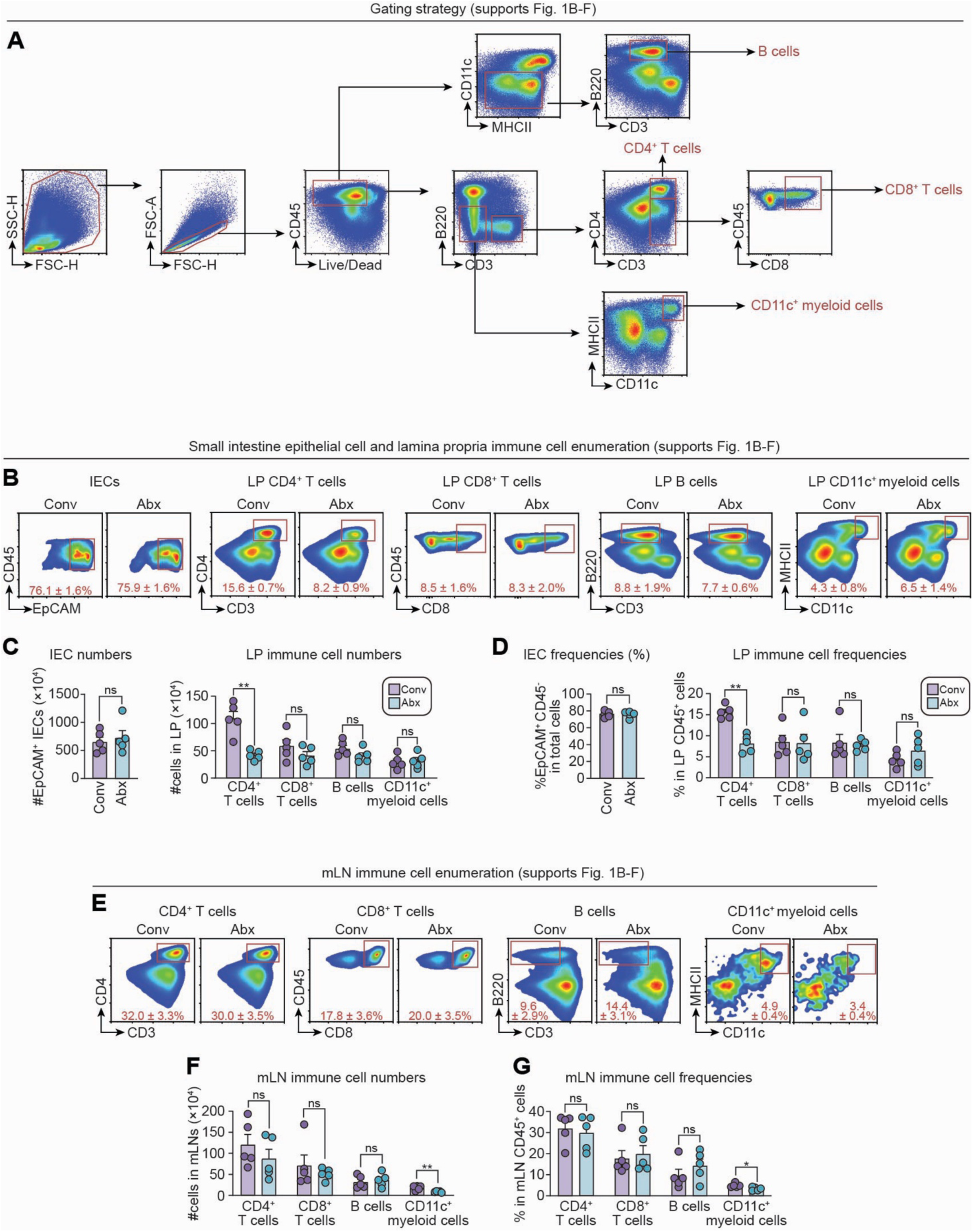
Flow cytometry gating strategy and quantification of epithelial and immune populations in the small intestine and mesenteric lymph nodes of conventional and antibiotic-treated mice (supports Figure 1). **(A)** Gating strategy for identifying immune populations in the small intestinal lamina propria and mesenteric lymph nodes. Cell subsets include CD4⁺ T cells (live CD45⁺B220^-^CD3⁺CD4⁺), CD8⁺ T cells (live CD45⁺B220⁻CD3⁺CD4⁻CD8⁺), B cells (live CD45⁺B220⁺CD3⁻CD11c⁻), and CD11c⁺ myeloid cells (live CD45⁺CD3⁻B220⁻CD11c⁺MHCII⁺). Representative flow cytometry plots are shown. **(B)** Flow cytometry analysis of IECs (live EpCAM^+^ CD45⁻) and lamina propria immune cells recovered from the small intestines of conventional and antibiotic-treated mice. Representative flow cytometry plots are shown. **(C)** Absolute numbers of IECs and LP immune cells from conventional and antibiotic-treated mice. **(D)** Relative frequencies of each cell population. % IECs is calculated as a percentage of the total cell population, and % LP immune cells is calculated as a percentage of total CD45^+^ cells. **(E)** Flow cytometry analysis of CD4⁺ T cells, CD8⁺ T cells, B cells, and CD11c⁺ myeloid cells recovered from mesenteric lymph nodes of conventional and antibiotic-treated mice. Representative flow cytometry plots are shown. **(F)** Absolute numbers of each mLN immune cell population from conventional and antibiotic-treated mice. **(G)** Relative frequencies of each mLN immune population (as a percentage of CD45⁺ cells) in conventional and antibiotic-treated mice. Conv, conventional; Abx, antibiotic-treated; IEC, intestinal epithelial cells; LP, lamina propria; mLN, mesenteric lymph nodes. n=5 mice per group. All data are representative of at least two independent experiments. Means ± SEM are plotted. \**p* < 0.05; \*\**p* < 0.01; ns, not significant by two-tailed Student’s *t*-test.

**Figure S3:**
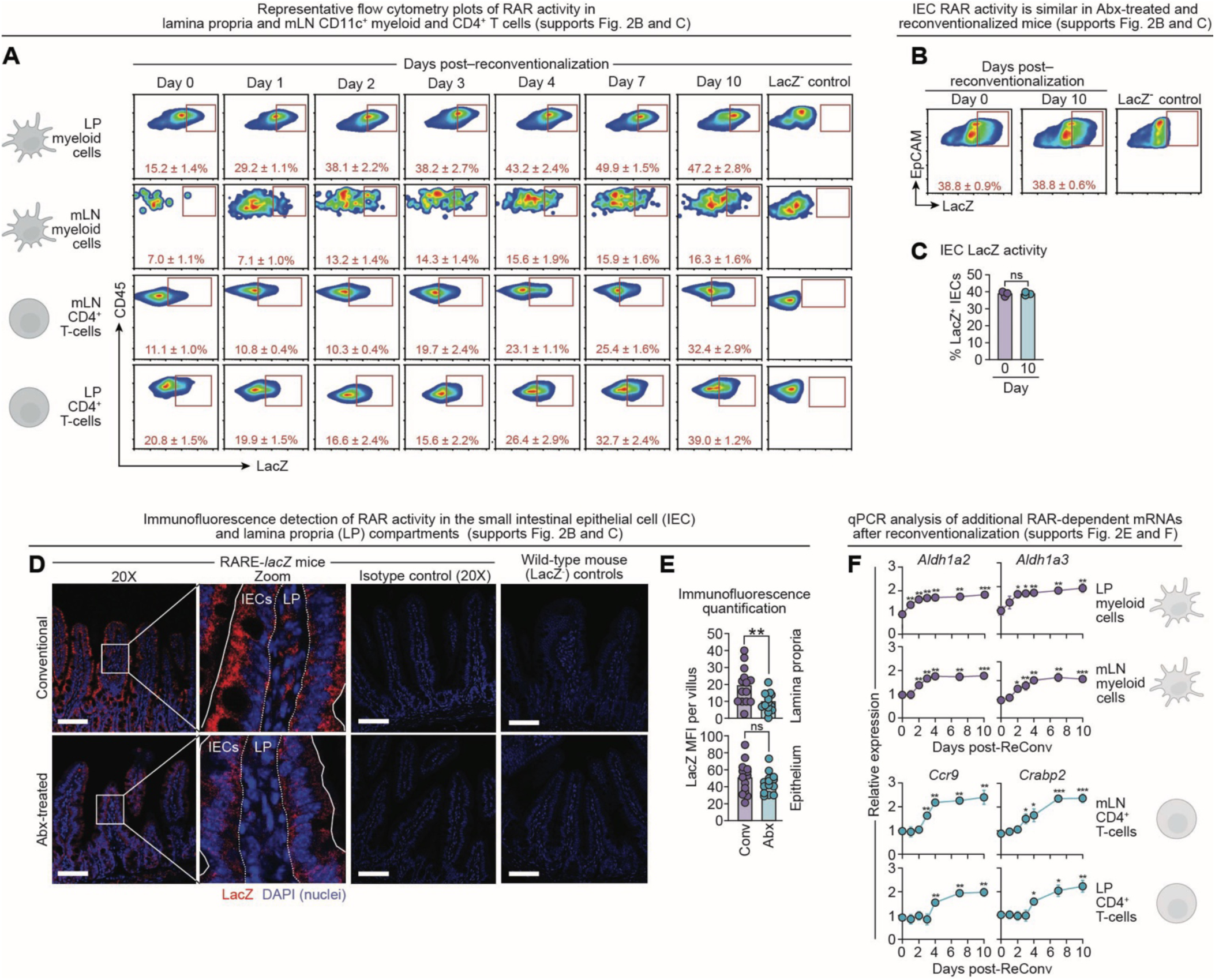
Time course of RAR activity and retinoid metabolism gene expression in CD11c⁺ myeloid cells and CD4⁺ T cells following microbiota reconstitution (supports Figure 2). **(A)** Representative flow cytometry plots showing LacZ⁺ (RAR⁺) cell frequencies at the indicated time points post-reconventionalization. Populations were gated as previously described: CD11c⁺ myeloid cells (live CD45⁺CD3⁻B220⁻CD11c⁺MHCII⁺) and CD4⁺ T cells (live CD45⁺B220⁻CD3⁺CD4⁺). LacZ activity was then plotted relative to CD45 expression. **(B)** Flow cytometry identification of RAR^+^ intestinal epithelial cells (IECs) at day 0 and 10 post-reconventionalization. IECs were gated as live EpCAM^+^CD45^-^. LacZ activity was then plotted relative to CD45 expression. Representative flow cytometry plots are shown. **(C)** Frequency of RAR⁺ IECs at day 0 and day 10 post-reconventionalization. n=3 mice per group per time point. **(D)** Immunofluorescence detection of LacZ in the small intestine of conventional and antibiotic-treated RARE*-lacZ* mice. Nuclei stained with DAPI. Scale bars: 100 μm (20X). Isotype and LacZ⁻ controls are shown. **(E)** Quantification of LacZ signal (MFI: mean fluorescence intensity) in the lamina propria and epithelial compartments. MFI of LacZ detection was determined for each villus across three fields per mouse. n=5 mice per group. **(F)** Time-course of RAR-dependent gene expression as measured by qPCR in LP CD11c^+^ myeloid cells, mLN CD11c^+^ myeloid cells, mLN CD4^+^ T cells, and LP CD4^+^ T cells. RAR, retinoic acid receptor; IEC, intestinal epithelial cells; Conv, conventional; Abx, antibiotic-treated; ReConv, reconventionalized; LP, lamina propria; mLN, mesenteric lymph node. n=3-5 mice per group. Means ± SEM are plotted. \**p* < 0.05; \*\**p* < 0.01; \*\*\**p* < 0.001; ns, not significant by two-tailed Student’s *t* test.

**Figure S4:**
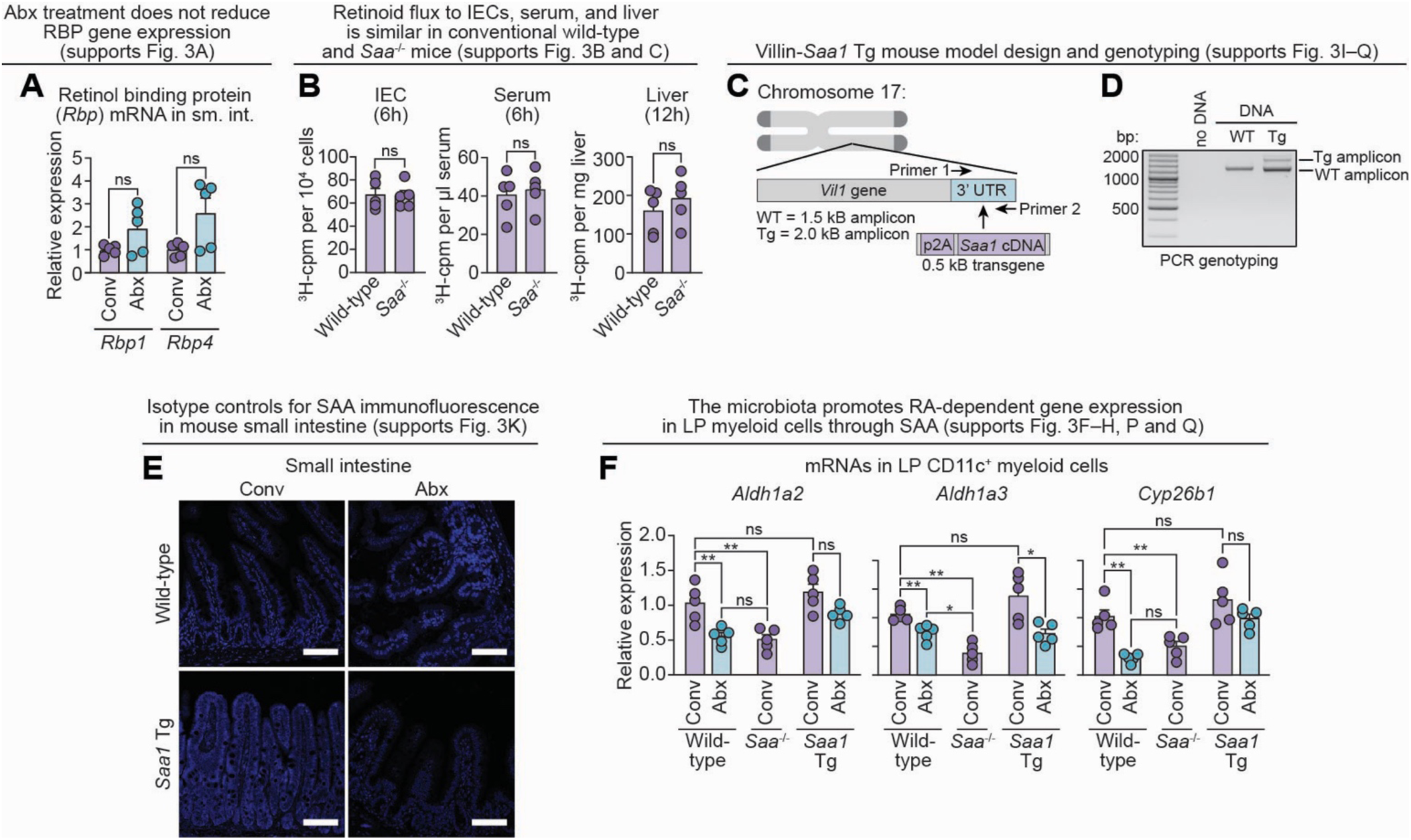
SAA-dependent regulation of intestinal retinol transport, systemic retinoid uptake, and myeloid cell retinoid metabolism (supports Figure 3). **(A)** qPCR analysis of *Rbp1* and *Rbp4* mRNAs in the small intestine of conventional and antibiotic-treated wild-type mice. *Rbp* genes encode a family of retinol-binding proteins (RBPs) that are distinct from the serum amyloid A family and that circulate in the bloodstream and within tissues.^43^ **(B)** ³H-retinoid uptake (normalized cpm per 10^4^ cells) in intestinal epithelial cells (IECs) 6 hours post-gavage. ³H-retinoid levels (normalized to volume) in serum at 6 hours post-gavage. ^3^H-retinoid levels (normalized to tissue mass) in the liver at 12 hours post-gavage. **(C)** Schematic of the Villin-*Saa1* transgene (*Saa1* Tg) construct. The *Saa1* transgene is inserted with a preceding p2A site into the 3′ UTR of the *Vil1* (Villin) gene to enable bicistronic transcription of *Vil1* and *Saa1* in intestinal epithelial cells.^45^ **(D)** PCR genotyping of wild-type and *Saa1* Tg mice. **(E)** Isotype controls for SAA immunofluorescence in ileal sections from conventional and antibiotic-treated wild-type and *Saa1* Tg mice. Scale bars = 100 μm. **(F)** qPCR analysis of RA-dependent genes in lamina propria CD11c⁺ myeloid cells from conventional and antibiotic-treated wild-type, conventional *Saa*⁻^/^⁻, and conventional and antibiotic-treated *Saa1* Tg mice. RBP, retinol-binding protein; IEC, intestinal epithelial cells; Tg, transgenic; Conv, conventional; Abx, antibiotic-treated; RA, retinoic acid; LP, lamina propria. n=3-5 mice per group. Means ± SEM are plotted. \*\**p* < 0.01; ns, not significant by two-tailed Student’s *t* test.

**Figure S5:**
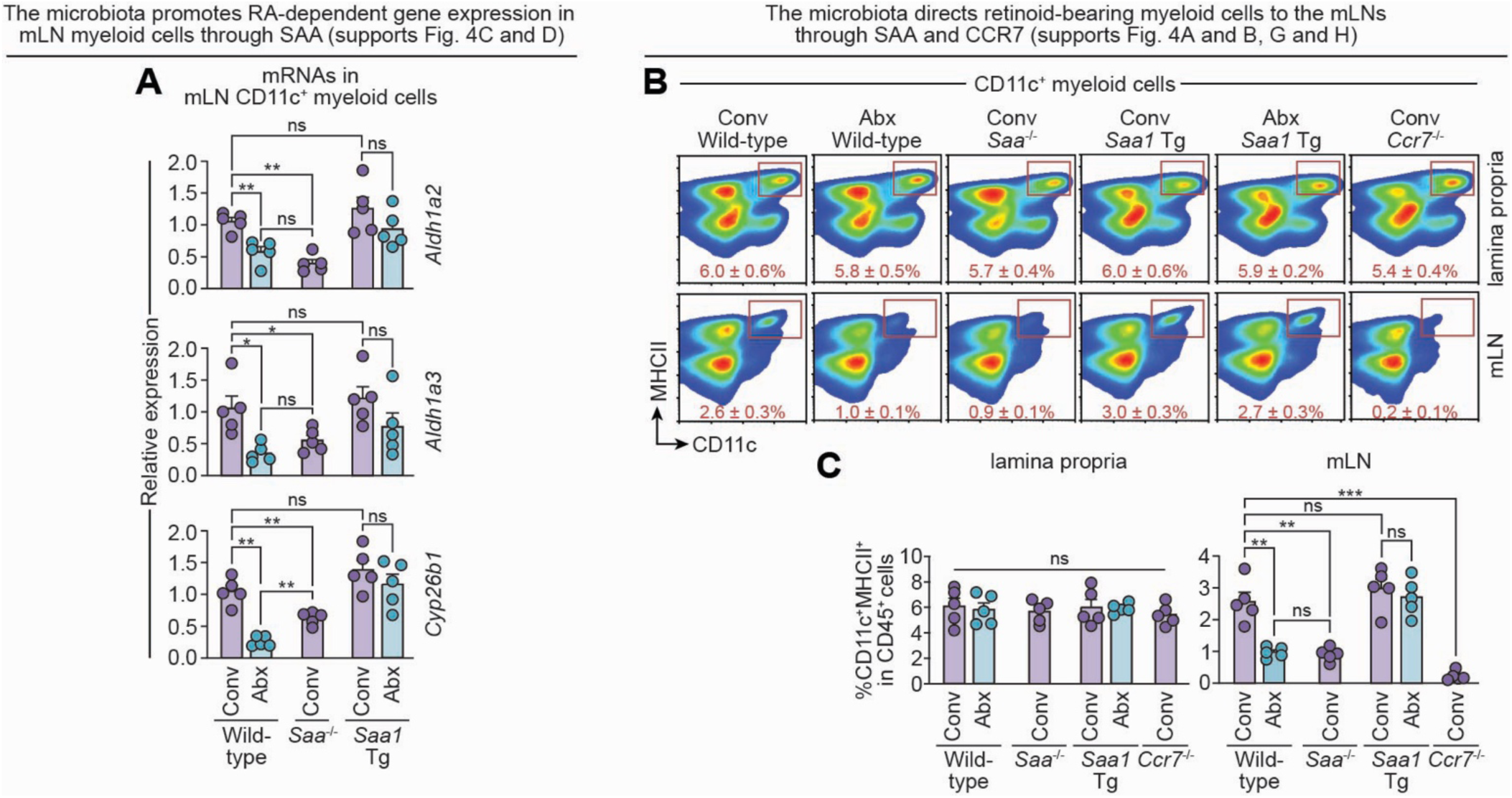
The microbiota directs retinoid-bearing myeloid cells to the mLNs (supports Figure 4). **(A)** qPCR quantification of RA-dependent mRNAs in magnetically enriched mLN CD11c⁺ myeloid cells from conventional and antibiotic-treated wild-type mice, conventional *Saa⁻^/^⁻* mice, and conventional and antibiotic-treated *Saa1* Tg mice. **(B)** CD11c⁺ myeloid cell enumeration by flow cytometry in the lamina propria and mesenteric lymph nodes of conventional and antibiotic-treated wild-type mice, conventional *Saa⁻^/^⁻* mice, conventional and antibiotic-treated *Saa1* Tg mice, and conventional *Ccr7*^-/-^ mice. Representative flow cytometry plots are shown. **(C)** Quantification of flow cytometry data in (B). RA, retinoic acid; mLN, mesenteric lymph nodes; Conv, conventional; Abx, antibiotic-treated. n=4-5 mice per group. All data are representative of at least two independent experiments. Means ± SEM are plotted. \**p* < 0.05; \*\**p* < 0.01; \*\*\**p* < 0.001; ns, not significant by two-tailed Student’s *t*-test

**Figure S6:**
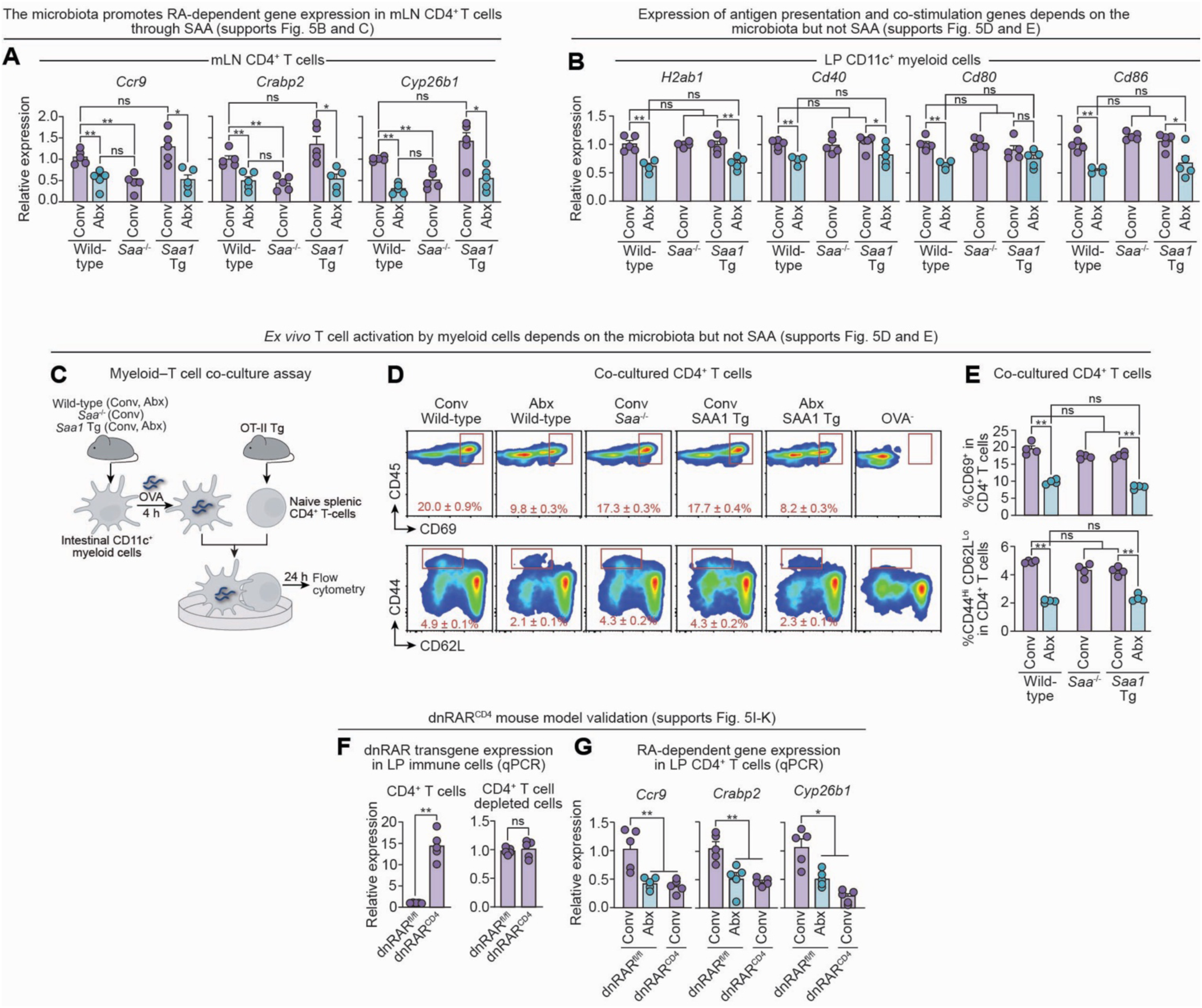
Role of the microbiota and SAA in myeloid cell antigen presentation, T cell activation, and gene expression (supports Figure 5). **(A)** qPCR quantification of RA-dependent mRNAs in magnetically enriched mLN CD4⁺ T cells from conventional and antibiotic-treated wild-type mice, conventional *Saa⁻^/^⁻* mice, and conventional and antibiotic-treated *Saa1* Tg mice. **(B)** qPCR quantification of mRNAs associated with antigen presentation and co-stimulatory molecules in LP CD11c⁺ myeloid cells isolated from the indicated mouse groups. **(C)** Strategy for myeloid–T cell co-culture assay. LP CD11c⁺ myeloid cells were isolated from conventional and antibiotic-treated wild-type mice, conventional *Saa⁻^/^⁻* mice, and conventional and antibiotic-treated *Saa1* Tg mice, pulsed with OVA_323-339_ peptide, and co-cultured with naïve splenic OT-II CD4⁺ T cells. T cells were then analyzed by flow cytometry. **(D)** CD4⁺ T cells co-cultured with myeloid cells as described in (C) were analyzed by flow cytometry. CD69 (top) and CD44 and CD62L (bottom) were used as markers of activation. **(E)** Quantification of the flow cytometry data in (D). **(F)** qPCR quantification of dnRAR transgene expression in CD4^+^ T cells recovered from the small intestine lamina propria of dnRAR^fl/fl^ and dnRAR^CD4^ mice (left). The same cell population depleted of CD4^+^ T cells by magnetic separation was also analyzed (right). **(G)** qPCR quantification of RA-dependent mRNAs in small intestinal CD4^+^ T cells from conventional and antibiotic-treated dnRAR^fl/fl^ and dnRAR^CD4^ mice. RA, retinoic acid; mLN, mesenteric lymph nodes; LP, lamina propria; Conv, conventional; Abx, antibiotic-treated; Tg, transgenic; OVA_323-339_, Ovalbumin peptide 323-339. n=4-5 mice per group. All data are representative of at least two independent experiments. Means ± SEM are plotted. \**p* < 0.05; \*\**p* < 0.01; ns, not significant by two-tailed Student’s *t*-test

**Figure S7:**
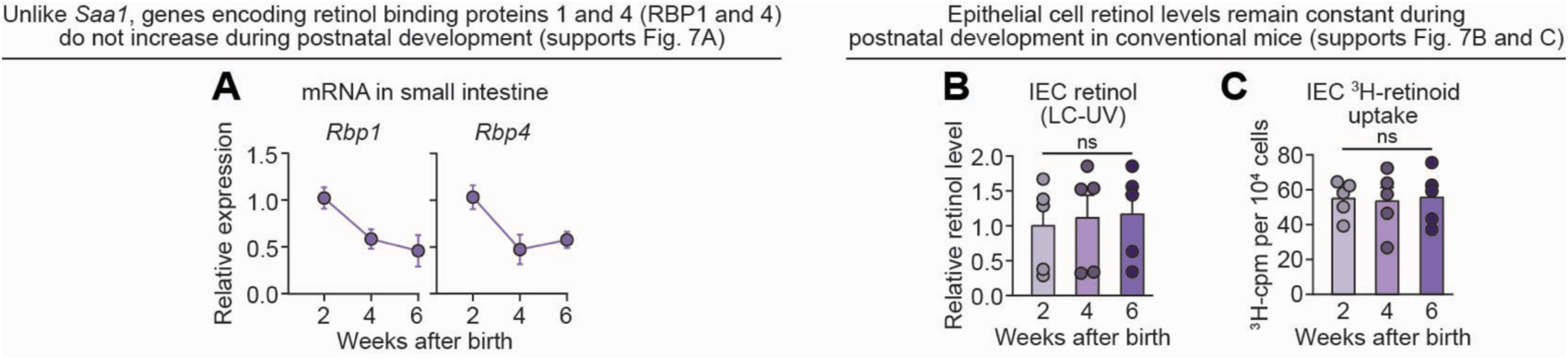
*Rbp* expression and retinoid uptake in intestinal epithelial cells across postnatal intestinal development (supports Figure 7). **(A)** qPCR analysis of *Rbp1*, and *Rbp4* mRNA expression in the small intestines of 2-, 4-, and 6-week-old conventional wild-type mice. n = 5 mice per group per timepoint. **(B)** Relative retinol content measured by LC-UV analysis of IECs isolated by EDTA treatment from 2-, 4-, and 6-week-old conventional wild-type mice. **(C)** ³H-retinoid levels in IECs 6 hours after gavage in 2-, 4-, and 6-week-old conventional wild-type mice. IEC, intestinal epithelial cells; RBP, retinol-binding protein. n=5 mice per group. Data are representative of two independent experiments. Means ± SEM are plotted. ns, not significant by two-tailed Student’s t-test.

## Experimental Model and Subject Details

### Mice

C57BL/6 wild-type, RARE*-lacZ*,^37^ *Saa^-/-^*,^42^ *MyD88*^ΔIEC^,^81^ *MyD88*^ΔCD11c^,^3^ *Rorc*^gfp/gfp^,^60,82^ *Rag1*^-/-^,^61^ *Stat3*^ΔIEC^,^3^ and dnRAR^CD4^ mice were bred and maintained under specific pathogen–free (SPF) conditions at the University of Texas Southwestern Medical Center. RARE*-lacZ*, *MyD88*^fl/fl^, ^83^ Villin-Cre,^45^ CD11c-Cre,^84^ CD4-Cre,^85,86^ *Rorc*^gfp/gfp^,^82^ *Stat3*^fl/fl^,^87^ *Ccr7^-/-^*,^88^ and OT-II transgenic mice were obtained from The Jackson Laboratory.^48^ *Saa*^-/-^ mice were generated by deletion of the *Saa* locus as previously described.^42^ dnRAR^fl/fl^ mice were obtained from the laboratory of Shanthini Sockanathan (Johns Hopkins University).^55^ All floxed strains were maintained as homozygotes; all Cre driver lines were maintained as heterozygotes. *MyD88*^ΔIEC^ mice were generated by crossing *MyD88*^fl/fl^ mice with Villin-Cre mice as previously described.^81^ *MyD88*^ΔCD11c^ mice were generated by crossing *MyD88*^fl/fl^ mice with CD11c-Cre mice as previously described.^3^ *Stat3*^ΔIEC^ mice were generated by crossing *Stat3*^fl/fl^ mice with Villin-Cre mice as previously described.^58^ dnRAR^CD4^ mice were generated by crossing dnRAR^fl/fl^ with CD4-Cre mice as previously described.^56^

Villin*-Saa1* transgenic (Tg) mice were generated in the UT Southwestern Transgenic Core. A p2A-linked *Saa1* cDNA was inserted into the 3′ untranslated region (UTR) of the endogenous *Vil1* (Villin) gene to enable epithelial-specific transgene expression (Fig. S4C,D). The self-cleaving p2A peptide was positioned upstream of *Saa1* to permit bicistronic transcription and independent translation of Villin and SAA1 proteins.^45^ CRISPR/Cas9- mediated homology-directed repair was used to insert the p2A-*Saa1* cassette. Guide RNAs targeting the *Vil1* 3′ UTR directed Cas9 cleavage, and a donor template containing the cassette flanked by homology arms was microinjected into zygotes. Founder mice were screened by PCR to confirm successful integration and were maintained on a C57BL/6J background.

Germ-free C57BL/6J mice were housed and bred in flexible film isolators at the UT Southwestern Gnotobiotic Facility. Unless otherwise specified, experiments were performed using male and female mice aged 6– 12 weeks. For postnatal developmental analyses, mice were evaluated at 2, 4, and 6 weeks of age. All animals were maintained on a 12-hour light/dark cycle with ad libitum access to standard chow and water. Mice were euthanized by isoflurane overdose, followed by cervical dislocation. All procedures were approved by the UT Southwestern Institutional Animal Care and Use Committee.

### Bacterial Strains

*Bacteroides thetaiotaomicron* (VPI-5482) was cultured anaerobically at 37°C in brain heart infusion (BHI) broth supplemented with hemin and vitamin K. *Enterococcus faecalis* (ATCC 29212) and *Escherichia coli* (MG1655) were grown aerobically at 37°C in BHI and Luria-Bertani (LB) broth, respectively; *E. coli* cultures were shaken during incubation. *Salmonella enterica* serovar Typhimurium (strain 1433) was cultured aerobically in LB broth at 37°C.

Segmented filamentous bacteria (SFB) were obtained from Andrew Gewirtz (Georgia State University) and propagated by monocolonizing germ-free mice. Cecal contents from these mice were harvested and used for subsequent monocolonization experiments.

## Method Details

### Isolation and flow cytometry analysis of immune cells from the lamina propria and mesenteric lymph nodes

Immune cells were isolated from the small intestinal lamina propria and mesenteric lymph nodes as previously described.^31^ For intestinal preparations, Peyer’s patches were excised, and the tissue was flushed with 10 mL ice-cold PBS, opened longitudinally, cut into ∼2 cm pieces, and washed three times by vortexing in 10 mL PBS with 5% FBS for 2 minutes each, using a multitube vortexer. Tissues were then incubated in 10 mL of HBSS containing 5% FBS and 10 mM EDTA at 37 °C for 15 minutes on a New Brunswick RM Innova 40 series platform shaker (at 225 rpm). Epithelial cells and intraepithelial lymphocytes were subsequently released by vortexing and removed by filtration through a 100 μm cell strainer. This EDTA incubation step was repeated once more to ensure complete removal of epithelial cells.

Remaining tissue was washed in PBS + 5% FBS, then transferred to 10 mL of digestion buffer [RPMI, 5% FBS, 0.2 mg/mL collagenase IV (Sigma), 0.15 mg/mL DNase I (Sigma), and 0.75 U/mL Dispase II (Sigma)]. Tissues were digested at 37°C for 45 minutes with agitation (225 rpm), vortexed for 2 minutes, and filtered through a 40 μm strainer. The digestion step was repeated once more on residual tissue to maximize immune cell yield.

Lamina propria cells were enriched by Percoll density centrifugation and collected from the 40–80% interphase, then washed twice with PBS + 3% FBS. Red blood cells were lysed using ACK buffer. Cells were used immediately for downstream applications.

For mesenteric lymph node (mLN) preparations, only lymph nodes draining the small intestine were excised. Nodes were placed in RPMI + 10% FBS, mechanically dissociated, and passed through a 40 μm filter. Red blood cells were removed using ACK buffer, and cells were immediately used.

For flow cytometry, single-cell suspensions were incubated with live/dead viability dye and anti-CD16/32 (Fc Block) for 10 minutes at 4 °C, followed by staining with fluorophore-conjugated antibodies (see Materials Checklist) for 25 minutes at 4 °C. After washing, samples were acquired on a NovoCyte 3005 flow cytometer and analyzed using NovoExpress software (ACEA Biosciences).

### Intestinal epithelial cell isolation

Intestinal epithelial cells (IECs) were isolated as previously described.^80^ Briefly, washed intestinal tissue was incubated in 10 mL of HBSS supplemented with 5% FBS and 10 mM EDTA at 4°C for 20 minutes. IECs were collected and immediately processed for downstream applications.

### Magnetic enrichment of CD11c^+^ myeloid cells and CD4^+^ T cells

Immune cells were isolated from the lamina propria and mLNs and subjected to magnetic enrichment using Miltenyi Biotec protocols. For CD11c⁺ myeloid cells, the CD11c MicroBeads Ultrapure kit was used according to the manufacturer’s instructions, with LS columns for magnetic separation. To maximize purity, myeloid cells underwent two consecutive rounds of enrichment. For CD4⁺ T cells, the Total CD4⁺ T Cell Isolation Kit was used per manufacturer’s instructions, with final magnetic separation also performed using LS columns.

### Antibiotic treatment and reconventionalization of wild-type mice

C57BL/6J wild-type mice housed under specific pathogen–free (SPF) conditions were considered conventional. To deplete the intestinal microbiota, mice received drinking water supplemented with 0.95 g/L Penicillin G sodium and 2 g/L Streptomycin sulfate for 10 days.^89,90^ Mice remained on antibiotic-containing water for the duration of experimental use and are referred to as antibiotic-treated. Microbiota depletion was confirmed by 16*S* qPCR analysis of fecal pellets.^58,89^ For reconventionalization, antibiotic-treated mice were switched to untreated water for 48 hours, then gavaged with 100 μL of a fecal slurry prepared from conventional donor mice. The slurry was generated by suspending 100 mg of freshly collected fecal pellets in 1 mL sterile PBS under anaerobic conditions, homogenizing for 10 min, and centrifuging at 1,000 × *g* for 10 min; the clarified supernatant was used for gavage. Mice receiving this treatment were designated reconventionalized.

### ^3^H-retinol flux assays

^3^H-retinol (11,12-[^3^H(N)]-retinol; Perkin Elmer) was diluted in corn oil and administered to mice by gavage (1 μCi per mouse in 100 μL of corn oil).^31^ For kinetic studies, mice were euthanized at 0 hours, 6 hours, and at 1, 2, 3, 4, 7, and 10 days post-gavage. Intestinal epithelial cells (IECs) were isolated using 10 mM EDTA chelation as previously described.^80^ Immune cells were isolated from the small intestine and mLNs as previously described, and CD11c⁺ and CD4⁺ T cells were enriched by magnetic separation.^31^

Cells were solubilized in 0.5 mL Biosol (National Diagnostics), mixed with 3.5 mL Bioscint scintillation cocktail, and radioactivity was measured using a liquid scintillation counter. Counts per minute (cpm) were normalized to the number of live cells, determined by viability dye exclusion. A cpm threshold of 250 was applied; values below this were considered undetectable.

Initial kinetic analysis identified peak ^3^H-retinoid uptake at distinct timepoints after tracer gavage for each cell type of interest: 6 hours for IECs, 1 day for LP CD11c⁺ cells, 2 days for mLN CD11c⁺ cells, 3 days for mLN CD4⁺ T cells, and 7 days for LP CD4⁺ T cells. These timepoints were used in subsequent retinoid flux studies.

To quantify systemic movement of ^3^H-retinoids, serum and liver uptake were assessed. For serum, whole blood was obtained 6 hours, 12 hours, and 24 hours post-gavage via cardiac puncture, collected in clot activator tubes, and allowed to clot at room temperature for 30 minutes. Samples were then centrifuged at 2000 × *g* for 10 minutes at 4°C. The resulting supernatant (serum) was mixed with 3.5 mL Bioscint scintillation cocktail, and radioactivity was measured using a liquid scintillation counter. Cpms were normalized to the volume of the serum. For the liver, a piece of liver tissue (left lobe) was excised 6 hours, 12 hours, and 24 hours post-gavage. After weighing the tissue, it was placed in 0.5 mL Biosol, homogenized, and allowed to incubate at 50°C for 4 hours. The samples were then centrifuged at 10000 × g for 10 minutes, after which the supernatant was mixed with 3.5 mL Bioscint scintillation cocktail, and radioactivity was measured using a liquid scintillation counter. Counts per minute (CPM) were normalized to the mass of the tissue.

### Liquid chromatography–ultraviolet spectroscopy (LC-UV) quantification of retinol in intestinal myeloid and epithelial cells

CD11c⁺ myeloid cells and intestinal epithelial cells (IECs) were isolated from small intestinal tissue as previously described. Following isolation, cells were pelleted, snap-frozen in liquid nitrogen, and stored at −80°C. Sample preparation for retinoid analysis was carried out under yellow-light conditions. Samples were hydrolyzed in 1 mL of 50 mM potassium hydroxide at 37 °C for 10 minutes. Retinoids were extracted by liquid-liquid extraction by adding 1 mL of hexane. The extraction process was repeated twice, and the resulting organic phases were pooled, dried under a stream of nitrogen, and resuspended in 100 µL of methanol.

All samples were analyzed using an Agilent 1290 Infinity II high-performance liquid chromatography (HPLC) system equipped with a UV detector. Retinol was identified by its characteristic retention time and absorbance at 325 nm. A 15 µL aliquot of each sample was injected onto an Agilent InfinityLab Poroshell 120 EC-C18 column (2.1 mm × 150 mm, 2.7 µm) maintained at 35 °C. The mobile phase consisted of solvent A (90% methanol, 10% water) and solvent B (100% methanol). The flow rate was 0.7 mL/min. The elution gradient is detailed in Supplementary Table 1. All measurements were performed under light-protected conditions to minimize photoisomerization and degradation.

To validate peak identity and retention time, a retinol standard (Cerilliant) was prepared in methanol and analyzed under identical chromatographic conditions. The retention time of the retinol peak in the standard was used to confirm the corresponding peak in biological samples.

Following extraction, the remaining aqueous phase was saved for protein quantification. Total protein concentration was determined using the Pierce^TM^ BCA assay (Thermo Scientific) according to the manufacturer’s instructions and was read out on an Agilent BioTek Cytation 5 cell imaging multimode reader. Retinol content in each sample was normalized to protein concentration. Normalized values were then scaled to the mean of control samples, which was set to 1.0, and all other values were expressed relative to this baseline.

### Quantitative real-time PCR (qPCR) of retinoid metabolism following microbial reconstitution

Antibiotic-treated mice were reconventionalized and sacrificed at 0 hours and at 1, 2, 3, 4, 7, and 10 days post-gavage. CD11c⁺ myeloid cells and CD4⁺ T cells were isolated from the small intestine and mesenteric lymph nodes by magnetic enrichment, as described above. Cell pellets were snap-frozen and stored at –80°C until RNA extraction.

Samples were lysed in RLT buffer (Qiagen), and total RNA was extracted using the RNeasy Plus Universal Mini Kit on a QIAcube automated platform. Complementary DNA (cDNA) was synthesized from purified RNA using M-MLV Reverse Transcriptase (Thermo Fisher Scientific). Quantitative real-time PCR (qPCR) was performed using TaqMan Universal PCR Master Mix (Thermo Fisher Scientific) on a QuantStudio 7 Flex Real-Time PCR System (Applied Biosystems). Transcript levels were quantified using the comparative Ct (ΔΔCt) method and normalized to 18*S* rRNA expression. The following transcripts were assessed: *Aldh1a2*, *Aldh1a3*, and *Cyp26b1* in myeloid cells; *Ccr9*, *Crabp2*, and *Cyp26b1* in CD4⁺ T cells.

For experiments involving other mouse models, transcript analysis in CD11c⁺ myeloid cells and CD4⁺ T cells from the small intestine and mesenteric lymph nodes was performed using the same protocol.

### qPCR for intestinal retinol carrier protein expression

For analysis of intestinal retinol carrier expression (i.e., *Saa*1, *Rbp1*, *Rbp4*), ileal tissue from the small intestine was collected and stored in RNAlater (Thermo Fisher Scientific) at –80°C until RNA extraction. RNA extraction, cDNA synthesis, and qPCR were performed as described above.

### RAR activity quantification following microbial reconstitution

Antibiotic-treated RARE*-lacZ* mice were reconventionalized and sacrificed at 0 hrs and at 1, 2, 3, 4, 7, and 10 days post-reconventionalization. Immune cells were isolated from the small intestine and mesenteric lymph nodes. Retinoic acid receptor (RAR) activity was assessed by quantifying β-galactosidase (LacZ) activity using the FluoReporter LacZ Flow Cytometry Kit (Thermo Fisher Scientific). Cells were loaded with 2 mM fluorescein di-V-galactoside (FDG) at 37°C for 1 min. FDG hydrolysis was halted by the addition of 1.8 mL ice-cold 1.5 µM propidium iodide. Cells were then stained with fluorophore-conjugated antibodies to identify immune cell subsets. Fluorescein release, indicative of LacZ enzymatic activity and RAR signaling, was measured by flow cytometry.

### Immunofluorescence assay

Mouse small intestinal tissue (ileum) was rinsed in PBS, fixed overnight at 4°C in 4% paraformaldehyde, and paraffin-embedded. Tissue sections were dewaxed with two xylene washes, then rehydrated through a graded ethanol series (100%, 95%, 70%, 50%, and water). Antigen retrieval was performed by boiling slides in 1X Tris-EDTA Buffer (pH 9.0 (Abcam)) for 20 minutes, followed by two PBS washes. Sections were blocked at room temperature for 1 hour in a 10% donkey serum blocking solution in PBS, then incubated overnight at 4°C with primary antibodies (anti-LacZ, 1:200; anti-SAA, 1:200) diluted in blocking buffer. AF647 anti-chicken and AF488 anti-goat antibodies were diluted 1:400 and applied to the slides for 1 hour at room temperature in the dark. After final washes, slides were mounted using DAPI-containing Fluoromount-G (Southern Biotechnology). For immunofluorescence imaging, 1X zoom images were captured with a 10X, 20X, and 40X oil-immersion lens on a Leica SP8 confocal microscope using Leica’s LAS X software. ImageJ software was used for the analysis of SAA1/2 and LacZ staining in IEC and LP fractions. Mean fluorescence intensity (MFI) of the relevant channel was quantified within the region of interest for each villus.

### RALDH activity quantification by Aldefluor assay

RALDH activity was measured using the ALDEFLUOR^TM^ assay kit (Stemcell Technologies) per the manufacturer’s instructions. The Aldefluor assay detects ALDH enzyme activity, which converts the non-fluorescent substrate BODIPY-aminoacetaldehyde (BAAA) to the fluorescent product BODIPY-aminoacetate (BAA). Cells with high RALDH activity retain greater amounts of BAA, which can be detected by flow cytometry.

Immune cells isolated from the small intestine and mesenteric lymph nodes were resuspended in 1 mL assay buffer and incubated with the Aldefluor reagent. To enable proper gating, half of each sample was treated with the ALDH inhibitor diethylaminobenzaldehyde (DEAB) as a negative control. Samples were incubated at 37°C for 40 minutes, washed, and stained with surface antibodies to identify immune subsets. Aldefluor-positive (RALDH-high) cells were quantified by flow cytometry in the green fluorescence channel (520-540 nm).

### Confirmation of the dnRAR^CD4^ mouse model

To validate CD4⁺ T cell–specific expression of the *dnRAR* transgene in *dnRAR^CD4^* mice, small intestinal immune cells were isolated and subjected to magnetic enrichment for CD4⁺ T cells. Both CD4⁺ T cell–enriched and CD4⁺ T cell–depleted fractions were collected. RNA was extracted from each fraction, reverse transcribed into cDNA, and transgene expression was assessed by qPCR to confirm dnRAR expression in the CD4⁺ T cell fraction and absence in the non-CD4⁺ fraction. To further confirm functional activity of the model, expression of RA-responsive genes (*Ccr9*, *Crabp2*, and *Cyp26b1*) was measured in intestinal CD4⁺ T cells from dnRAR^fl/fl^ and dnRAR^CD4^ mice.

### Monocolonization of germ-free mice

Germ-free mice were orally inoculated with either *Bacteroides thetaiotaomicron* (VPI-5482), *Enterococcus faecalis* (ATCC29212), *Escherichia coli* (MG1655), each prepared at 2×10⁹ CFU in log-phase culture, or with 1×10⁹ CFU of log-phase *Salmonella enterica* serovar Typhimurium (strain 1433). Colonization was performed by gavage, and animals were euthanized three days post-inoculation. For segmented filamentous bacteria (SFB) monocolonization, cecal contents from SFB-monocolonized donor mice (kindly provided by Andrew Gewirtz, Georgia State University) were suspended in 1 mL of sterile PBS, briefly centrifuged at 1000 × *g* to remove debris. 100 μL of the clarified supernatant was administered by gavage. These mice were sacrificed three days after colonization.

### SFB colonization and depletion in conventional mice

To assess segmented filamentous bacteria (SFB) colonization, fecal pellets were collected and genomic DNA was extracted as described above. Quantitative PCR (qPCR) was performed using SFB-specific 16*S* rRNA primers (forward: 5′-GACGCTGAGGCATGAGAGCAT-3′; reverse: 5′-GACGGCACGGATTGTTATTCA-3′).^91^ Colonization status was determined by comparing amplification signals between SFB-positive and SFB-negative mice.

To generate SFB-positive mice, SFB-negative animals (obtained from Jackson Laboratories and confirmed by qPCR) were co-housed with SFB-colonized cagemates (obtained from Taconic Farms and confirmed by qPCR) for 2 weeks, and monitored for SFB acquisition via fecal qPCR.^5,58^ SFB depletion was achieved by providing drinking water containing 0.95 g/L Penicillin G sodium and 2 g/L streptomycin sulfate for 10 days^58^. Depletion was confirmed by absence of detectable SFB signal by qPCR.

### OT-II CD4⁺ T cell ^3^H-retinol uptake in co-culture with intestinal myeloid cells

CD11c⁺ myeloid cells were isolated from the small intestinal lamina propria of wild-type mice and plated in 96-well U-bottom plates (5 × 10⁴ cells/well) in 200 μL of complete RPMI. Cells were incubated for 4 hours with 1 nCi ^3^H-retinol and with or without 200 ng OVA₃₂₃–₃₂₉ peptide. Following incubation, wells were washed to remove excess peptide and unincorporated tracer. Naïve OT-II CD4⁺ T cells (2.5 × 10⁵ cells/well), isolated from OT-II spleens by magnetic enrichment (Miltenyi Naïve CD4^+^ T cell isolation kit), were added at a 1:5 myeloid:T cell ratio and co-cultured for 24 hours. Cells were then washed with PBS three times to remove dead cells and supernatant, solubilized in 0.5 mL Biosol (National Diagnostics), mixed with 3.5 mL Bioscint scintillation cocktail, and analyzed on a liquid scintillation counter.

### OT-II CD4⁺ T cell activation by intestinal myeloid cells

CD11c⁺ myeloid cells were isolated by magnetic enrichment from the small intestinal lamina propria of conventional and antibiotic-treated wild-type mice, conventional *Saa^-/-^* mice, and conventional and antibiotic-treated Villin-*Saa1* transgenic mice. Cells were plated in 96-well U-bottom plates (5 × 10⁴ cells/well) in 200 μL of complete RPMI with or without 200 ng OVA₃₂₃–₃₂₉ peptide and incubated for 4 hours. Wells were washed to remove excess peptide, and naïve OT-II CD4⁺ T cells (2.5 × 10⁵ cells/well), isolated from OT-II spleens by magnetic enrichment as above, were added at a 1:5 myeloid:T cell ratio. After 24 hours, T cell activation was assessed by flow cytometry using CD69, CD44, and CD62L staining.

### Bulk RNA sequencing and transcriptomic analysis of mLN CD4^+^ T cells

RNA was isolated from magnetically enriched CD4⁺ T cells from the mesenteric lymph nodes of wild-type and *Saa⁻^/^⁻* mice, using RLT buffer (Qiagen), and the RNeasy Plus Universal Mini Kit on a QIAcube automated platform as described above. RNA integrity was verified using an Agilent 2100 Bioanalyzer. Libraries were prepared using the Takara SMART-Seq mRNA kit and the Kapa HyperPlus library preparation kit. Samples were sequenced on the Illumina NovaSeq 6000 platform, generating paired-end 150 bp reads.

Raw reads were trimmed for adapters and low-quality bases using Trim Galore. Reads were mapped to the mouse reference genome (mm39) using STAR (v2.7.10b) with gene annotations from the UCSC Genome Browser and NCBI RefSeq.^92^ Prior to alignment, library quality was assessed by mapping to mouse transcript and ribosomal RNA sequences using BWA (v0.7.17).^93^ SAMtools (v1.16.1) was used to sort alignments.^94^ Gene-level counts were generated with HTSeq, and differential expression was assessed using DESeq2 in Bioconductor.^95–97^ Differential expression was defined by an adjusted *p*-value < 0.05. KEGG pathways and GO annotations (NCBI gene2go) were queried for enrichment using Fisher’s exact test in R.^98^

### Developmental retinoid flux and RAR activity assays

To quantify retinol in intestinal CD11c^+^ myeloid cells, these cells were isolated from conventional wild-type mice, conventional *Saa⁻^/^⁻* mice, and germ-free wild-type mice at 2, 4, and 6 weeks of age. Retinol levels were measured by liquid chromatography with UV detection (LC-UV). This measurement was also performed for isolated IECs.

For assessment of ^3^H-retinoid uptake, conventional wild-type mice, conventional *Saa⁻^/^⁻* mice, and germ-free wild-type mice aged 2, 4, and 6 weeks were gavaged with ^3^H-retinol. Three days later—based on previously established uptake kinetics—mice were sacrificed and mLN CD4⁺ T cells were isolated for scintillation counting.

RAR activity in mLN CD4⁺ T cells was assessed in conventional and antibiotic-treated RARE*-lacZ* reporter mice at 2, 4, and 6 weeks of age by flow cytometry, as previously described. To generate antibiotic-treated 2-week-old mice, breeding cages were maintained on antibiotic-supplemented water for 10 days starting at birth, after which the offspring were designated as antibiotic-treated.

To assess RA-dependent gene expression in T cells, mLN CD4^+^ T cells were isolated from conventional wild-type mice, conventional *Saa⁻^/^⁻* mice, and germ-free wild-type mice aged 2, 4, and 6 weeks, as previously described. RNA was extracted from each fraction, reverse-transcribed into cDNA, and RA-dependent gene expression was quantified by qPCR.

Intestinal T cell infiltration was quantified in conventional wild-type mice, conventional *Saa⁻^/^⁻* mice, and germ-free wild-type mice aged 2, 4, and 6 weeks by flow cytometry of lamina propria immune cells.

### Statistical analysis

Specific statistical methods, definitions of significance, and the number of replicates are detailed in the corresponding figure legends. Data are presented as mean ± standard error of the mean (SEM). The number of replicates reflects biologically independent experiments conducted on separate days.

Sample sizes were not predetermined using formal power calculations. Experiments were not blinded, and no formal randomization protocol was applied; however, mice were assigned to groups without bias, and samples were processed in a non-systematic order. Animals that died before the experimental endpoint were excluded from analysis.

All statistical analyses were performed using GraphPad Prism. Differences between two groups were evaluated using two-tailed Student’s *t* tests. Additional statistical tests, if applied, are specified in the figure legends.

## Supplementary tables

**Table 1.**

| Time (min) | Mobile A (%) | Mobile B (%) |
| --- | --- | --- |
| 0.0 | 100 | 0 |
| 2.5 | 0 | 100 |
| 8.5 | 0 | 100 |
| 8.6 | 100 | 0 |
| 10.0 | 100 | 0 |

## Key Resources Table

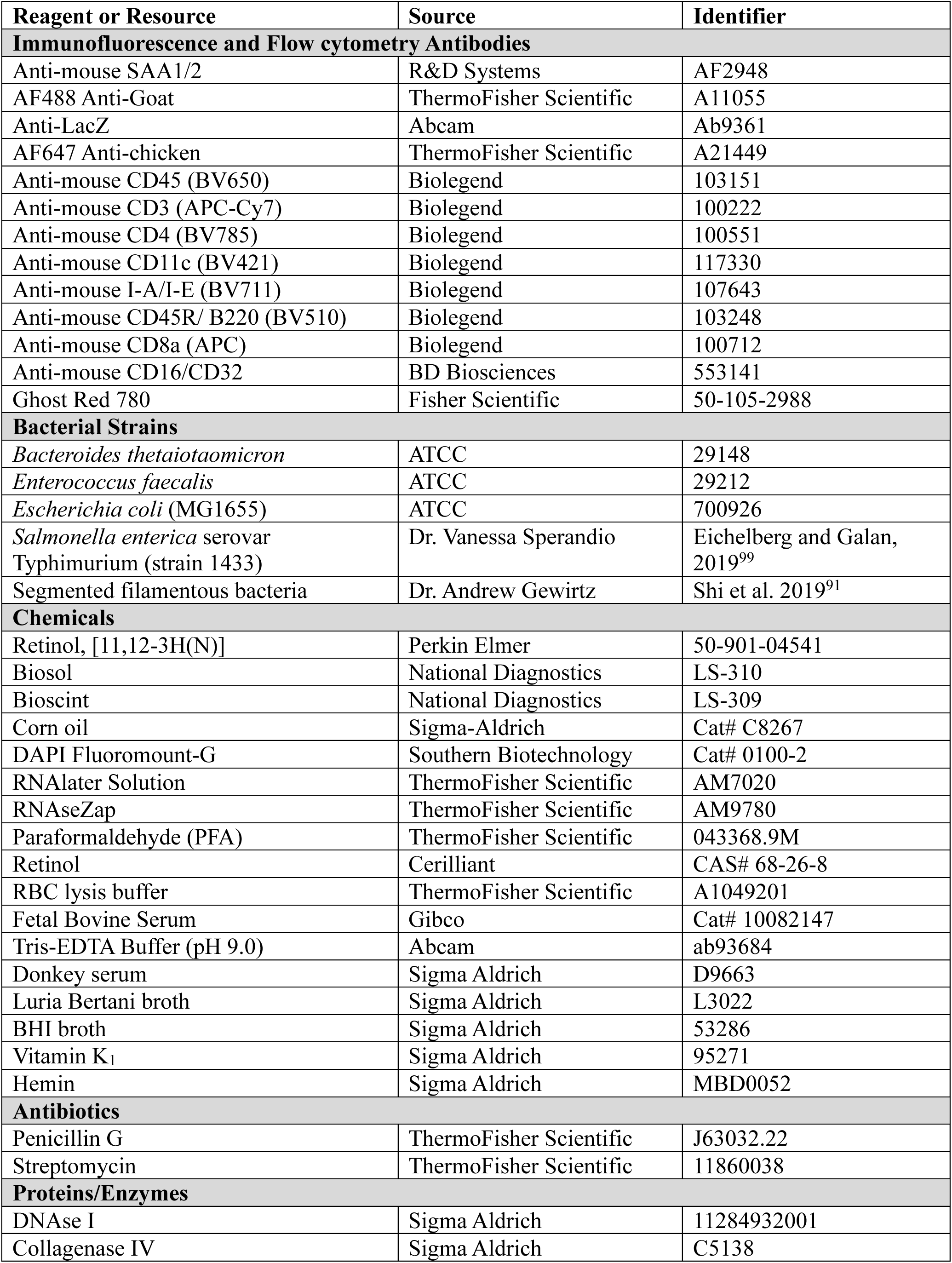
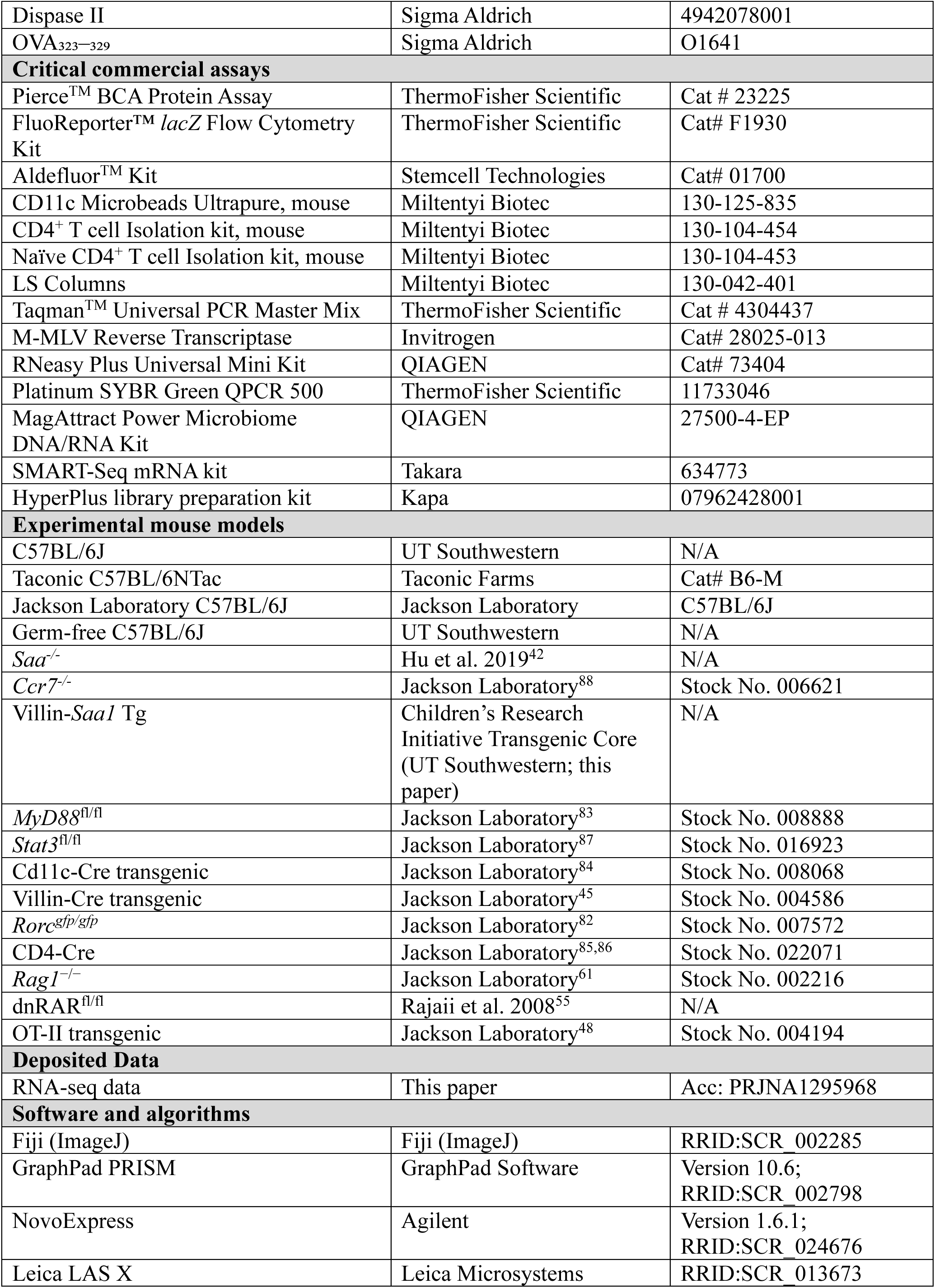
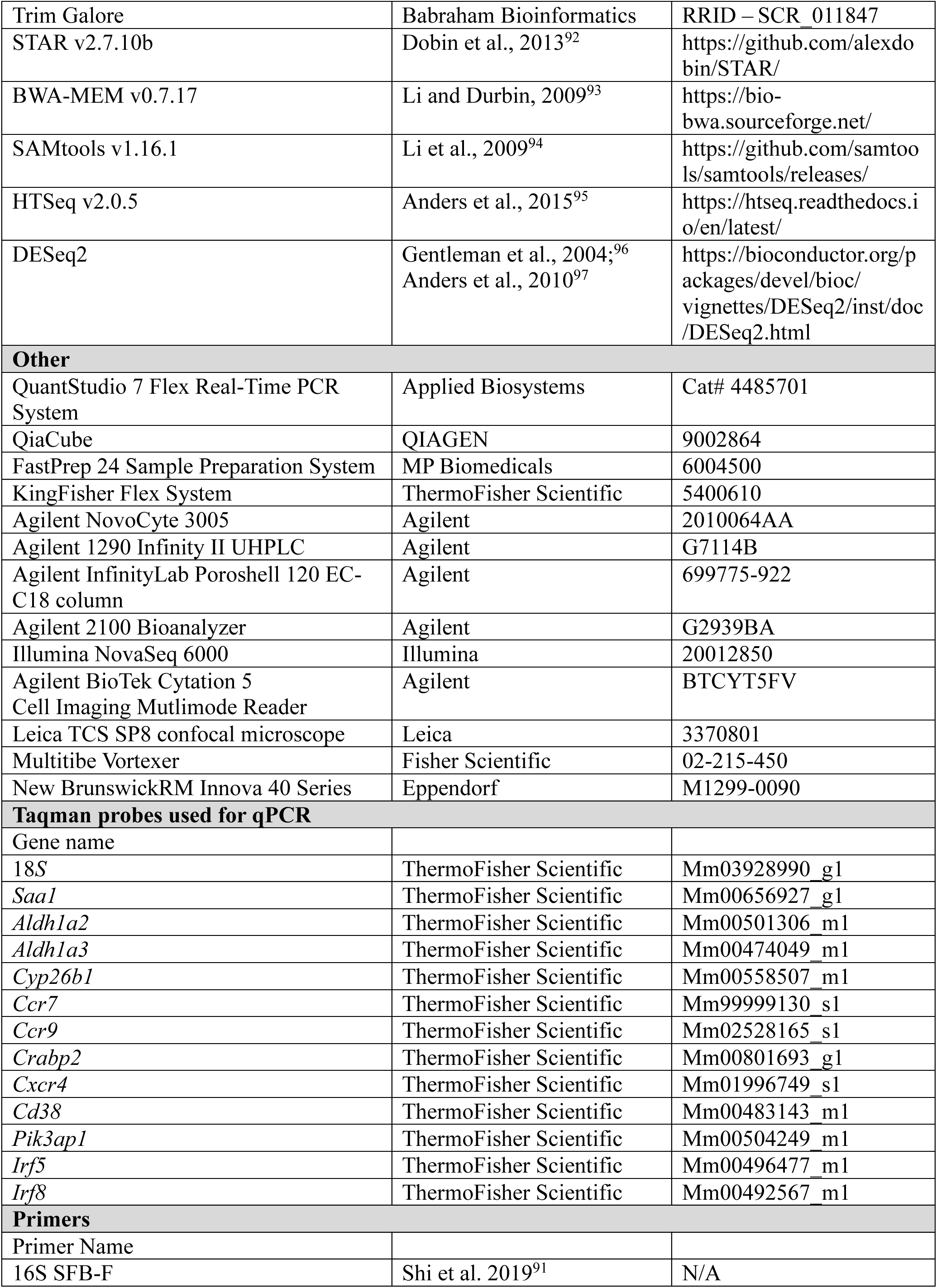
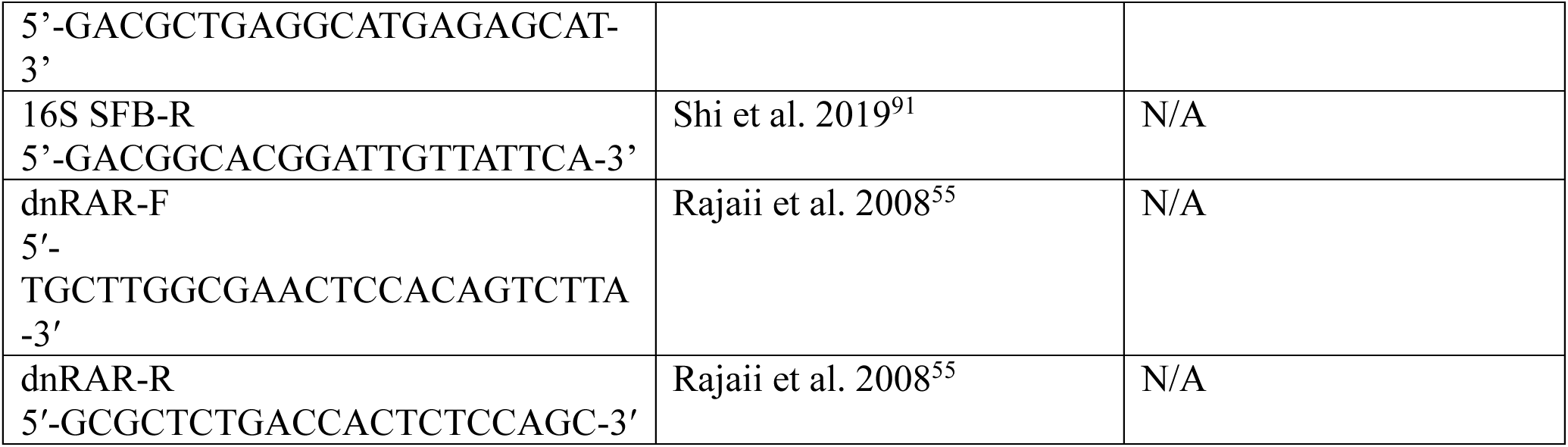

## Data Availability

RNA-seq data are available from the Gene Expression Omnibus (GEO) repository under accession no. PRJNA1295968. All other data are available in the main text or the supplementary materials.

## Contact for Reagent and Resource Sharing

Further information and requests for resources and reagents should be directed to and will be fulfilled by the Lead Contact, Lora V. Hooper. The dnRAR mouse line was obtained from Shanthini Sockanathan and cannot be distributed to other groups without permission from Dr. Sockanathan.

## Acknowledgements

We thank Matthew Johnson and Devyn Slaven for assistance with mouse experiments, and Dr. Shanthini Sockanathan (Johns Hopkins University) for providing the dnRAR^fl/fl^ mice. This work was supported by NIH grants R01 DK070855 (L.V.H.), R01 CA231303 (A.Y.K.), P01 AI179406 (A.Y.K. and L.V.H.); Welch Foundation Grant I-1874 (L.V.H.); the Walter M. and Helen D. Bader Center for Research on Arthritis and Autoimmune Diseases (L.V.H.); the University of Texas Southwestern Medical Center and Children’s Health Cellular and Immuno Therapeutics Program (A.Y.K.); and the Howard Hughes Medical Institute (L.V.H.). T.S. was supported by NIH T32 AI007520. Components of some figures were generated using BioRender.

## Author Contributions

T.S., A.Y.K., L.V.H. designed the research. T.S., C.D., K.A.R., S.L.M., A.J., C.L.B., B.H., G.V., J.N.L., C.L., C.A. performed the research. X.Z., J.G.M. provided experimental materials. T.S., A.J., G.V, J.K., C.A., P.R. analyzed the data. T.S. and L.V.H. wrote the paper.

